# Sequence-based drug design using transformers

**DOI:** 10.1101/2023.11.27.568880

**Authors:** Shengyu Zhang, Donghui Huo, Robert I. Horne, Yumeng Qi, Sebastian Pujalte Ojeda, Aixia Yan, Michele Vendruscolo

## Abstract

Protein-ligand interactions play central roles in biological processes and are of key importance in drug design. Deep learning-based approaches are emerging as cost-effective alternatives to high-throughput experimental methods for the screening of large libraries of ligands. Here, to predict the binding affinity between proteins and small molecules, we introduce Ligand-Transformer, a deep learning framework based on the AlphaFold2 transformer architecture. We applied Ligand-Transformer to screen inhibitors targeting the mutant EGFR^LTC^ kinase, identifying compounds with low nanomolar potency. We then used this approach to predict the conformational population shifts induced by ABL kinase inhibitors. To show the applicability of Ligand-Transformer to disordered proteins, we explored the binding of small molecules to the Alzheimer’s Aβ peptide, identifying compounds that delayed its aggregation. Overall, Ligand-Transformer illustrates the potential of transformers in accurately predicting the interactions of small molecules with both ordered and disordered proteins, thus uncovering molecular mechanisms and facilitating the initial steps in drug discovery.

## Introduction

Recent reports are revealing that AlphaFold2 and related approaches exhibit capabilities beyond their original function of predicting protein structures^1–11^. To build on these results to develop protein-ligand binding prediction methods, we report Ligand-Transformer, a deep learning approach designed to model protein-ligand complexes (**Figure S1**). Ligand-Transformer takes as input the protein sequence and the graph representation of the ligand, and predicts the binding affinity of the complex. The approach also generates distance matrices that predict the inter-residue distances of the protein, inter-atomic distances of the ligand, and atom-residue distances for the complex.

Ligand-Transformer utilizes the transformer framework of AlphaFold2^1^ to generate protein representations from their sequences, and the Graph Multi-View Pre-training (GraphMVP) framework^12^ to generate ligand representations. Instead of using the final predicted protein structure, we leverage the intermediate outputs of AlphaFold2. For the ligands, during pre-training, GraphMVP injects the knowledge of 3D molecular geometry into a 2D molecular graph encoder, allowing downstream tasks to benefit from the implicit 3D geometric prior. By leveraging the prior knowledge encoded in these high-dimensional representations, we capture the structural features of protein-ligand interactions, resulting in an accurate modelling of the bound states. The representations are further processed by the main structure of Ligand-Transformer, which consists of three parts (**Figure S1**). The first part is the feature encoders to re-process the representations of proteins and ligands. The second part is the cross-modal attention network to exchange the information between representations of the protein and ligand. The third part is composed by two downstream predictors, the first head for affinity predictions and the second head for distance predictions.

### Performance comparison against state-of-the-art affinity prediction methods

We conducted a performance comparison of Ligand-Transformer against other affinity prediction methods^13–15^ utilizing the PDBbind2020 dataset (**Table S1**). The results indicate that Ligand-Transformer achieves comparably better correlations with experimentally-measured values when compared to the baseline methods.

### Identification of EGFR^LTC^ ligands

We illustrate the potential of Ligand-Transformer for the identification of initial hits in drug discovery pipelines by screening ligands targeting EGFR^LTC^, a mutant form of the EFGR kinase. EGFR is a key target in cancer therapy due to its role in cell growth^16^. As mutations in EGFR, such as L858R/T790M/C797S (LTC), can lead to resistance against all current EGFR inhibitors, there is a need for novel drugs to target this triple mutant^17^. We first collected a dataset, called EGFR^LTC^-290 (**Supplementary Data 2**), consisting of 290 existing inhibitors with their measured half maximal inhibitory concentration (IC_50_) values, along with annotations indicating whether they are allosteric or orthosteric (see Methods). We found that Model B (see Supplementary Information) could predict the binding affinity with a Pearson’s correlation coefficient (R) value of 0.57 (**Figure 1a**). To fine-tune the model on this specific dataset to achieve higher accuracy, we randomly split the EGFR^LTC^-290 dataset into ten parts, and conducted a ten-fold cross validation to evaluate the performance and obtain an ensemble of fine-tuned models (Model FT1 to FT10, see Methods). The value of R of this test increased to 0.88 after fine tuning.

**Figure 1.**
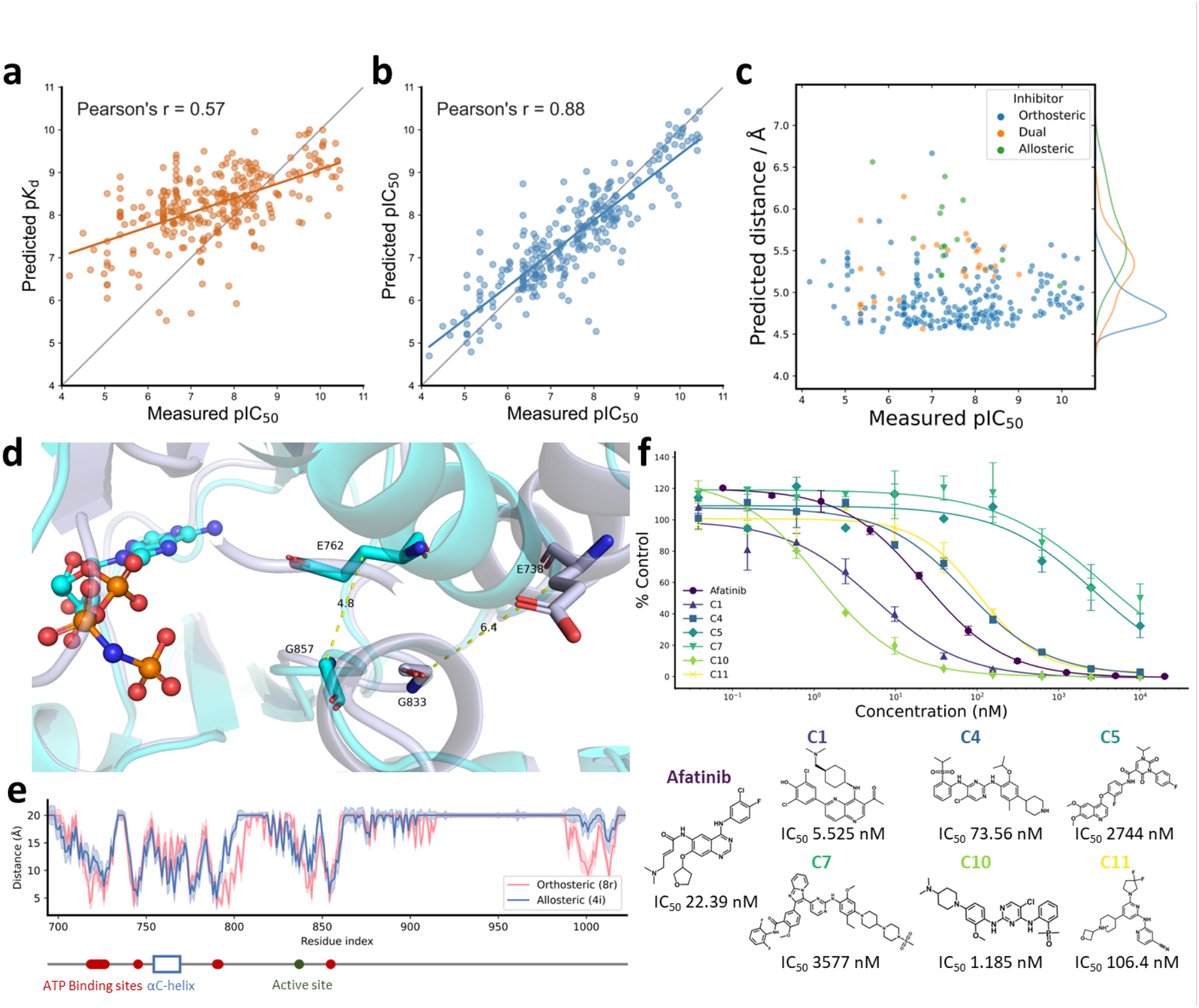
Identification of ligands targeting EGFR^LTC^. **(a**, **b)** Scatter plot of the correlation between the experimental pIC_50_ values and the binding parameters (p*K*_d_ and pIC_50_ values) using Ligand-Transformer for complexes in the EGFR^LTC^-290 dataset; all binding parameters are in molar units: **(a)** predicted p*K*_d_ values without transfer learning (base model), and (**b**) predicted pIC_50_ values post ten-fold validation with transfer learning. (**c**) Scatter plot representing the distribution of predicted distances between E762 and G857 of EGFR^LTC^ in various ligand-binding states. The y-axis represents the predicted distance between the E762 Cβ and G857 Cα atoms, while the x-axis depicts the experimental pIC_50_ values of the ligands. The data points are coloured based on their reported binding sites in the literatures: orthosteric (blue), allosteric (green), and dual (orange). The kernel density estimations (KDE) plot for the distribution of predicted distances are indicated adjacent to y coordinates. (**d)** Comparative visualization of EGFR kinase domain crystal structures in the active (PDD 2ITX) and inactive (PDBID: 2GS7) states. The αC-helix is depicted as a ribbon structure in light blue and gray, representing the active and inactive states, respectively. The AMP-PNP ligand bound to PDB 2ITX is visualized as a stick-and-ball structure. Residues E762 and G857 of EGFR (corresponding to residues E738 and G833 in PDB 2GS7) are shown as sticks. Nitrogen, oxygen, and phosphorus atoms are colored blue, red, and orange, respectively. (**e)** Line graph of the predicted binding modes of an orthosteric inhibitor (8r , colored in red) and an allosteric inhibitor (4i ^43^, colored in blue) with EGFR^LTC^. The line graph illustrates the predicted minimal distances between each residue and the ligand. The error bars represent the predicted confidence scores pMAE (see **Algorithm 8** in Supplementary Iinformation). The bottom part shows sequence annotation of EGFR based on UniProt P00533. ATP binding sites are shown in red dots, active site is shown in green dot, and the region of αC-helix is shown in blue box. (**f)** In vitro kinase inhibition assays for selected compounds (C1 to C11), depicted as a percentage of activity relative to a DMSO control. The graph presents the mean ± standard deviation, with each ligand tested in duplicate. The corresponding IC_50_ values are presented for each compound.

We identified notable differences in the binding modes of the predicted orthosteric and allosteric inhibitors. Statistical analysis using the *t*-test revealed no significant differences in activity distribution between the orthosteric and allosteric groups (p=0.66) and the dual group (p=0.39) (**Figure 1b**). However, the predicted distance distributions exhibited significant variations (p<10^-21^ for allosteric, p=0.0025 for dual). The predicted distance between residues E762 and G857 was significantly greater when binding to allosteric inhibitors compared to orthosteric inhibitors. This distance can serve as an indicator of the DFG-in and DFG-out states of the αC-helix (**Figure 1c**), consistent with previous studies^18,19^. Specifically, the DFG-in state of the αC-helix represents the active conformation, while allosteric inhibitors tend to bind to DFG-out state of the αC-helix, which is the inactive conformation. Moreover, the residue-wise distance between the kinase and ligand indicates that the binding sites of allosteric inhibitors are situated in closer proximity to the αC-helix region rather than the ATP-binding site (**Figure 1d**).

We then used the predicted affinities and distances obtained from Ligand-Transformer to screen a subset of TargetMol containing 9,090 compounds in stock (see Methods and **Supplementary Data 3**). We selected candidates with the criteria that they are predicted to have high binding affinity by all of the 11 models (see Supporting Information). Finally, we obtained 12 candidates (**Extended Data Table 1**) with predicted IC_50_ between 1 to 100 nM). These candidates included a compound previously identified as effective inhibitor targeting EGFR^LTC^ (known as brigatinib) with a reported IC_50_ of 1-38 nM^20^. To our knowledge, among the remaining 11 compounds, C1, C4, C5, C10, and C11 have not been reported to target EGFR, while C7 is known to target EGFR but not to EGFR^LTC^.

We then tested experimentally the inhibitory potency of these 11 candidates, and found six active compounds. Out of these six active compounds, three fell within the predicted IC50 range of 1 to 100 nM, with two of them, C1 and C10, exhibiting high potency, with IC_50_ values of 5.5 and 1.2 nM, respectively (**Figure 1f**). C1 is a naphthyridine derivative, which is a non-traditional scaffold for EGFR inhibitors, it is not a known tyrosine kinase inhibitor. Based on our predictions, the naphthyridine moiety may be capable of competing with the purine ring of ATP and forming hydrogen bonds with the backbone of M793, thereby stabilizing the ligand within EGFR pocket. C10 and C4 show Tanimoto similarities (based on topological fingerprints in RDKit^21^) of 0.97 and 0.54, respectively, to brigatinib. These compounds exhibit a high degree of pharmacophore overlap, particularly the aniline-pyrimidine scaffold. This structural motif may compete with the purine ring of ATP, occupying the EGFR pocket. Additionally, the pyrimidine moiety can form hydrogen bonds with backbones of M793 and P794, thereby enhancing its binding affinity. C5, C7, and C11 exhibit inhibitory activity against EGFR^LTC^ in the range of 100 nM ∼ 10 μM, indicating relatively weak inhibitory potency. C11 is a pyridine compounds, which consists of only two aromatic rings. From a structural perspective, it represents a novel EGFR inhibitor. C5 is a pyrimidine-2,4-dione derivative and features a quinoline ring, which is a classic structural motif commonly found in EGFR inhibitors. On the other hand, C7 has a complex structure and a relatively high molecular weight, which does not align with the characteristics of ATP-competitive inhibitors targeting EGFR.

### Conformational selectivity of ABL kinase inhibitors

Kinases are dynamic, interconverting between different conformational states^22^. Monitoring these transitions and characterizing the conformational states that a kinase populates has proven to be a significant challenge^23^. Kinases typically maintain highly conserved active states to ensure the accurate positioning of essential catalytic regions^18^. In contrast, their inactive states can be distinct among different kinases^24^. Examining how small molecules selectively bind individual kinases could be instrumental in developing inhibitors specifically targeting these unique inactive states^24^. Here, we investigated whether Ligand-Transformer can be used to predict the ligand binding-induced conformational population shift of the ABL kinase, which plays a pivotal role in several signalling pathways, governing crucial cellular processes such as growth, survival, invasion, adhesion, and migration^25,26^.

ABL is reported to have three major conformational states, one active state (A) and two inactive states (I_1_ and I_2_)^23^. To investigate the ability of Ligand-Transformer to capture the change of the conformation ensemble of ABL after binding, we collected 12 inhibitors of this kinase, each with a predominant state of ABL when bound, as determined by nuclear magnetic resonancy (NMR) spectroscopy^23^. A total of 60 possible structures of ABL were obtained from the PDB (entries 6XR6, 6XR7, and 6XRG), with 20 structures corresponding to each of the three states A, I_1_ and I_2_ (**Figure 2a**). We then used the predicted distances between the residues of ABL as constraints to reweight the conformational ensemble consisting of the 60 structures and calculate the population of each conformational state (see Methods). Upon binding, the predicted predominant states are in accordance with experimental measurements for 11 of 12 compounds (**Figure 2b**). Furthermore, when grouping the inhibitors by their labelled predominant state, we found that each group had a significantly higher population within the corresponding state compared to the other two groups.

**Figure 2.**
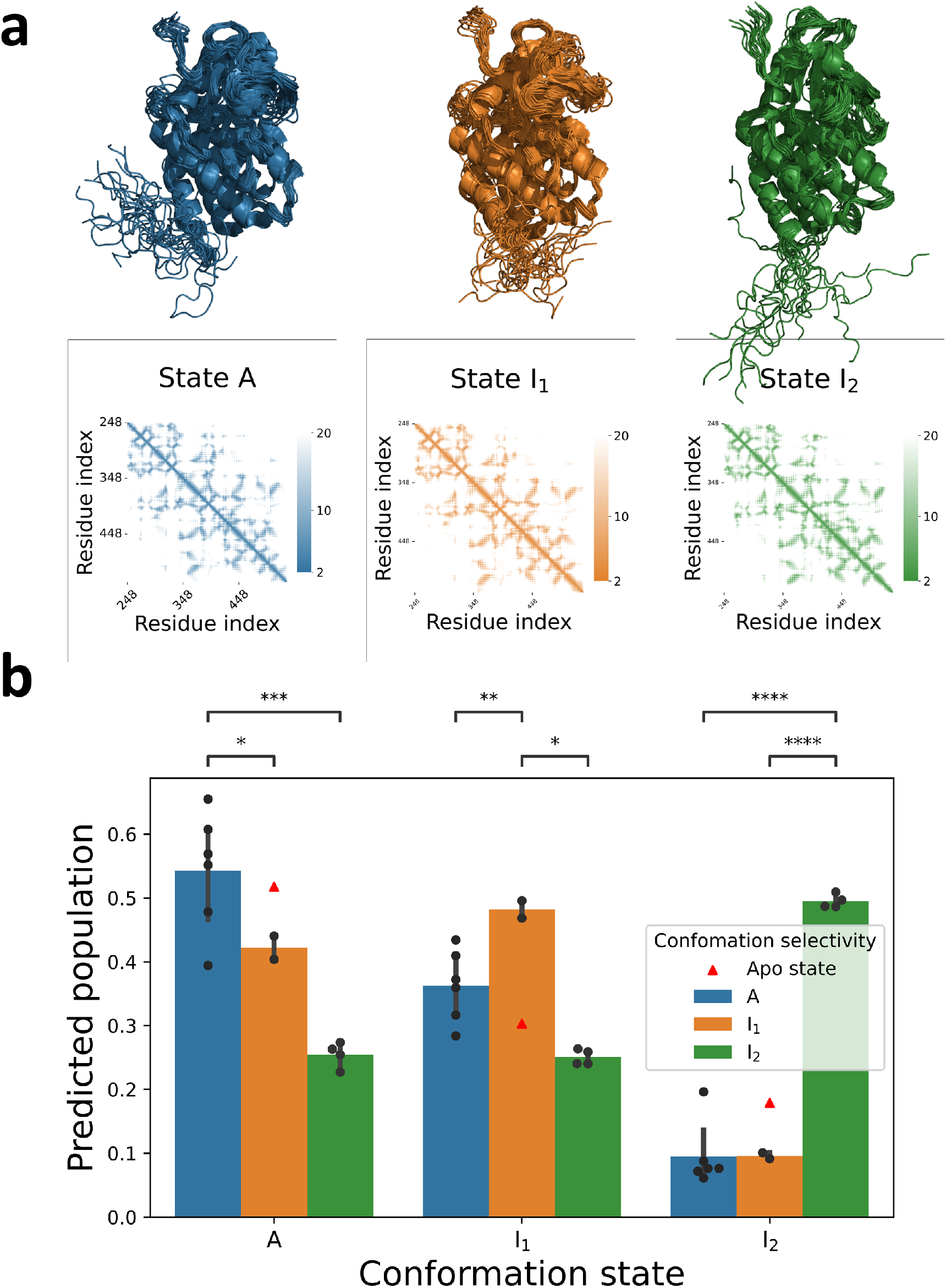
Conformational selectivity of inhibitors targeting ABL kinase. **(a)** Average distance maps and structural ensembles for ABL kinase states A (blue), I_1_ (orange), and I_2_ (green), corresponding to PDBs 6XR6, 6XR7, and 6XRG respectively. (**b)** Conformation selectivity prediction of different inhibitors. Twelve molecules were divided into three groups based on their conformational selectivity as determined in the literature^23^: Group A (blue, n=6), Group I_1_ (orange, n=2), and Group I_2_ (green, n=4). The grouped bar graph illustrates the differences in predicted population of binding state conformations of ABL when interacting with inhibitors from different groups. The error bars represent the standard deviation within each group. The red triangle represents the population of ABL conformations predicted by AlphaFold2 in the apo state. The Welch’s *t*-test was used to assess statistical significance, denoted as *p<0.05, **p<0.01, ***p<0.001, and ****p<0.0001. Population estimations are derived from predicted distance matrices (see Methods for details of calculations). The predicted populations of ABL state bound with each individual ligand are listed in **Extended Data Table 2**

We conducted a detailed exploration of the predicted distance distributions that illustrate the distinct conformation ensembles of ABL when bound with various inhibitors. Our analysis (**Extended Data Figure 1**) focuses on the spatial positioning of the αC-helix (residues 299 to 311), the phosphate-binding loop (P-loop, residues 267 to 275), and the activation loop (A-loop, residues 400 to 424) of ABL. These elements are pivotal in differentiating the three states of ABL^23^. We chose four representative residue pairs, selected for their high F-statistics indicating their effectiveness in distinguishing between the three states, enabled us to investigate the distances between critical protein components.

**Extended Data Figure 2** illustrates that the distance between V308 and F401 represents the distance between the αC-helix and the DFG motif (comprising residues D400, F401, and G402, at the start of the activation loop). The increased distance in states A, I_1_, and I_2_ is notable, reflecting the DFG-out conformation in states I_1_ and I_2_, and the DFG-in state in state A^23^. The distance between residues V275 and F401 represents the gap between the DFG motif and the P-loop (**Extended Data Figure 3**). This distance is significantly reduced in state I_2_, which adopts a P-loop stretched conformation, as opposed to the P-loop kinked conformation in states A and I_1_^23^. Predictions from Ligand-Transformer corroborate this trend. The distance between residues H380 and G402 represents the span from the active site to the activation loop (**Extended Data Figure 4**). In state I_2_, which adopts an A-loop closed conformation^23^, this distance is larger than in the other states, and the measured structures show that the distance in state I_1_ is slightly shorter than in state A. In state I_2_, the A-loop-closed conformation results in a shorter distance between L403 and L406, indicating a more compact activation loop (**Extended Data Figure 5**).

**Figure 3.**
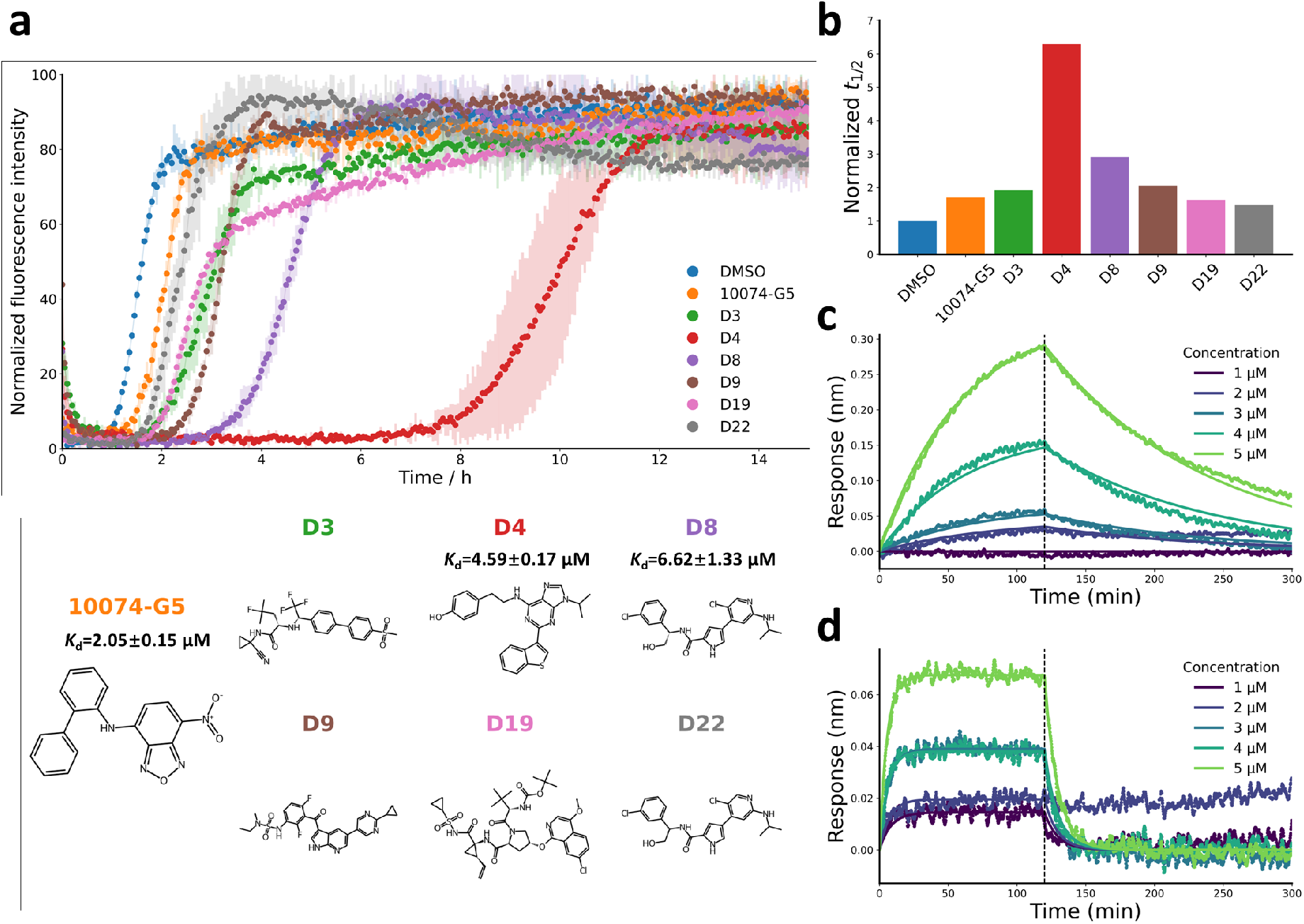
Identification of compounds targeting Aβ42 in its monomeric disordered state. **(a)** Kinetic fluorescence profiles of a 2 μM solution of Aβ42 at 37 °C, in the presence of 1% DMSO alone (blue), or in the presence of 20 μM of the predicted compounds (each represented by a different color). Error bars represent the mean ± standard deviation of two replicates. **(b)** Bar graph of the normalized half-time (*t*_1/2_) of Aβ42 aggregation in the presence of each ligand, relative to the DMSO control. **(c, d)** Biolayer interferometry (BLI) measurements showing the binding kinetics of compounds D4 (**c**) and D8 (**d**) to Aβ42-functionalized surfaces at different concentrations. Baseline drift were corrected, and global fitting to a one-phase association and dissociation model provided the kinetic rates. These fits were constrained to the same *k*_on_ and *k*_off_ values across all curves.

All of these observations of distance distribution differences are recapitulated in the predictions of Ligand-Transformer. Specifically, when ABL in the holo state is bound with an inhibitor selective to a specific state, it tends to favor a distance distribution pattern that aligns with the measured distribution of that particular state (**Extended Figures 2**-**5**). Moreover, the predicted distance distributions from Ligand-Transformer exhibit a wide range with multiple peaks, differing from the apo state predictions by AlphaFold2. This suggests that Ligand-Transformer can provide information about the structural ensembles of a protein-ligand complex.

### Identification of ligands of disordered proteins

Aβ is a disordered peptide that forms toxic aggregates in Alzheimer’s disease^27,28^. It has been suggested that small molecules could bind and stabilise the monomeric state of Aβ, thus slowing down its aggregation process^29^. A proof-of-concept for small molecules with this mechanism of action, known as pharmacological chaperones, has been provided by tafamidis, a drug approved for the treatment of certain types of transthyretin amyloidosis^30^. However, since the monomeric state of Aβ is disordered, it lacks stable binding pockets, making it challenging to apply structure-based drug design methods^31^. While several studies have made progress in developing methods for designing binders to disordered proteins, these approaches often on molecular dynamics simulations, thus requiring substantial computational resources^32–34^. To test whether Ligand-Transformer could offer a faster alternative computational screening method for disordered protein targets, we used it to predict the affinity of small molecules for the monomeric state of the 42-residue form (Aβ42) of Aβ to find aggregation inhibitors. From the ZINC-Cayman library containing 8,112 molecules (**Supplementary Data 4**), we thus selected 29 molecules. These were chosen based on their high predicted activity and favorable Central Nervous System Multiparameter Optimization (CNS MPO) scores, as detailed in the Methods section. In addition to these 29 molecules, we also included 8 purchasable enantiomers derived from this subset for experimental testing.

We studied the formation of Aβ42 fibrils in vitro with and without the presence of the selected compounds. In the absence of these compounds, Aβ42 aggregated with a half-time of approximately 2 h under our buffer conditions. For 6 of the 38 tested compounds (**Extended Data Table 3**), which were introduced at a concentration of 20 μM (i.e. 10 molar equivalents, Meq), we noticed a delay in Aβ42 aggregation (half-time > 3 h, **Figure 3a,b**). Among these compounds, D4 stood out as the most effective in delaying Aβ42 aggregation, extending the half-time to 6.7 times that of the control condition.

Biolayer interferometry (BLI, see Methods) was used to measure the binding affinity between small molecules and monomeric Aβ42. Firstly, N-terminally biotinylated Aβ42 monomer was immobilized on the surface of super streptavidin sensor tips (see Methods) due to the strong affinity between streptavidin and biotin. Then the tips were exposed to small molecules at five different concentrations (1 to 5 µM). The figures below represent the binding curves of the five small molecules with double subtraction (both the baseline drift and non-specific binding have been corrected). The binding response observed in 10074-G5, D4, and D8 turned out to be dose-dependent, indicative of binding (**Extended Figure 6a** and **Figure 3c,d**, respectively), while D3 and D9 showed weak and non-systematic binding response, indicative of weak binding. We globally fit the binding curves under 1:1 binding model to a simple one-step association and dissociation equations, which give a shared single association (*k*_on_) and dissociation (*k*_off_) rates, thus a shared dissociation constant (*K*_d_) in each molecule. For 10074-G5, the *k*_on_ was 3.83 × 10^4^ M^−1^s^−1^ and *k*_off_ was 7.85 × 10^-2^ M^−1^s^−1^, which gives a *K*_d_ value of 2.05 ± 0.15 μM. For D4, the *k*_on_ was 1.84 × 10^3^ M^−1^s^−1^ and *k*_off_ was 8.45 × 10^-3^ M^−1^s^−1^, which gives a *K*_d_ value of 4.59 ± 0.17 μM. For D8, the *k*_on_ was 1.59 × 10^4^ M^−1^s^−1^ and *k*_off_ was 1.06 × 10^-1^ M^−1^s^−1^, which gives a *K*_d_ value of 6.62 ± 1.33 μM. The binding results of 10074-G5, D4, and D9 are comparable to other-small molecule interactions with disordered proteins^35^.

## Discussion

We have proposed a sequence-based method of predicting the binding affinity of ligands for proteins. This approach also predicts the corresponding binding mode, represented as distance matrices between the proteins and the ligands. Through comparisons with other baseline models^13–15^ and ablation experiments, we observed that Ligand-Transformer performs well in affinity predictions. This result can be attributed to the protein and molecular representations provided by pre-trained AlphaFold2^1^ and GraphMVP^12^, as well as the structural information learned from the distance matrices of protein-ligand complexes during the training.

Because conformational fluctuations are crucial in determining protein behavior^36–38^, predicting conformational ensembles remains a crucial element for the development of next-generation structural prediction algorithms^39^. In this context, Ligand-Transformer offers an approach towards understanding the impact of ligand binding on conformational ensembles using predicted distance matrices. Specifically, the rational design of allosteric modulators of kinases, particularly type-4 inhibitors^24^ and activators^40^, often encounters challenges with the traditional lock-and-key paradigm. Our methodology proposes a potential solution, enabling the design of ligands that modulate kinase activity by influencing the population distribution across various conformational states.

We have also reported six compounds that inhibit Aβ42 aggregation by acting as pharmacological chaperones, highlighting the potential utility of our approach both for ordered and for disordered protein targets. Notably, by not relying on structures as input, and leveraging the capability of AlphaFold2 to encode information on the conformational space of disordered proteins^11^, Ligand-Transformer shows promise in addressing the challenges associated with designing strategies for disordered targets, which are still considered largely undruggable.

## Materials and Methods

### Protein representations generated by AlphaFold2

We utilized AlphaFold2, a pre-trained model to extract protein representations^1^. The protein representation consists of three key components. Firstly, we generated the single sequence representation 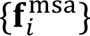, where 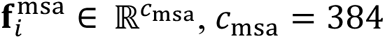, and 𝑖 ∈ {1 . . . 𝑁_res_}. This representation is derived by linearly projecting the first row of the multiple sequence alignment (MSA) representation. The MSA representation is the output of the final layer of the Evoformer and serves as the input to the structure module in AlphaFold2^1^. Secondly, we established pair representations 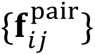 for each pair of residues 𝑖 and 𝑗, where 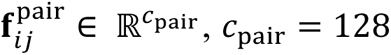. The pair representations 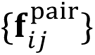 are also outputs of the Evoformer and the input to the structure module in AlphaFold2 and are used in predicting inter-residue distances in AlphaFold2. Lastly, the structure representation 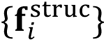 was obtained from the final layer of the structure module in AlphaFold2, where 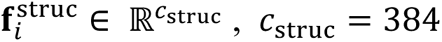, and 𝑖 ∈ {1 . . . 𝑁_res_} . This structure representation is employed in AlphaFold2 for predicting side-chain dihedral angles and model confidence prediction. The MSA searching was conducted by MMseqs2 (default setting) on BFD/MGnify and Uniclust30 (2021_03). No structural template was fed during the prediction. Model 1.1.1 of AlphaFold2 was used for the inference, and no structural template was used.

### Ligand representations generated by GraphMVP

For obtaining a stereospecific molecular geometry representation, we used GraphMVP, a pre-trained molecule encoder that transforms 2D ligands into graph representations^12^. GraphMVP is trained via self-supervised learning (SSL) using auxiliary tasks. Here, both the input atoms and chemical bonds underwent a one-hot encoding process before being input into GraphMVP. The outputs generated include the atom representation, denoted as 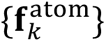, where 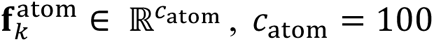, and 𝑘 ∈ {1 . . . 𝑁_atom_}, and the bond representation, represented as 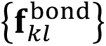, where 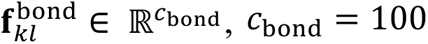, for each chemical bond formed between atom 𝑘 and atom 𝑙.

## Datasets

### PDBbind2020-subset

We utilized the publicly accessible PDBbind v2020 dataset^20^, which provides the structures of 19,443 protein-ligand complexes along with their experimentally measured binding parameters. We eliminated structures with more than one ligand and those whose ligand structures could not be read by RDKit (http://www.rdkit.org/). Additionally, to enhance the training efficiency, we initially limited the dataset to complexes where the ligand atom count was 128 or fewer, and the protein sequence length was restricted to 384 amino acids or fewer. For sequences exceeding 384 amino acids in length, we applied a truncation strategy (see Supplementary Information) by eliminating domains that were distant from the binding site. This resulted in a subset dataset (PDBbind2020-subset) consisting of 13,420 data points. We subsequently randomly divided the dataset into training, validation, and testing sets, containing 10,375, 640, and 936 data points respectively (see **Supplementary Data 1**). During model training, we initially conducted warm-up training with 4,480 complexes (PDBbind_4k) that only included single chains with lengths less than 384. Following this, we proceeded to train with the complete training set (PDBbind_10k) of 10,375 data points (see Supplementary Information for training details).

For activity data, the PDBbind dataset includes dissociation constant (*K*_d_), half maximal inhibitory concentration (IC_50_), and inhibition constant (*K*_i_) measurements. As prediction labels, we took the negative logarithm of these values (in molar units), resulting in the p*K*_d_, pIC_50_ and p*K*_i_ values, respectively. We considered *K*_i_ and *K*_d_ measurements without distinction to train Ligand-Transformer to predict *K*_d_ values. The IC_50_ data are assay-specific and only comparable under certain conditions. Based on literature research^41^, augmenting mixed public IC_50_ data with public *K*_i_ data does not deteriorate the quality of the mixed IC_50_ data if the *K*_i_ is corrected by an offset. Therefore, we did not explicitly distinguish among the three types of activity data. Instead, we corrected all IC_50_ values by dividing them by a factor 2.3 That is, we added 0.35 log units to all pIC_50_ values to obtain their p*K*_d_ equivalents for use as prediction labels. Ligand-Transformer can thus predict *K*_d_, IC_50_ and *K*_i_ values.

For the distance data, the ground truth distances are computed from the PDB files using Biopython As the distance prediction has a lower limit of 1 Å and an upper limit of 20 Å, all labels are truncated within this range when calculating the absolute error between the predicted distances and true distances.

### EGFR^LTC^-290

We collected 400 inhibitors targeting the L858R/T790M/C797S triple-mutant EGFR from ChEMBL^42^ and SciFinder (as of November 2022). These inhibitors were obtained from about 90 literature sources, and all data were double-checked with the original publications to ensure some important information, e.g., tested with human protein, and L858R/T790M/C797S triple-mutant EGFR form. For duplicates we retained the one with higher bioactivity, resulting in a final set of 290 inhibitors with IC_50_ values ranging from 0.08 nM to 150 μM. We classified the inhibitors into orthosteric inhibitors, allosteric inhibitors, or those that bind to both sites, based on information from their originally published literature.

### TargetMol library for screening EGFR^LTC^ inhibitors

The library used for virtual screening for EGFR^LTC^ inhibitors in this study contained a total of 9,090 compounds, all sourced from TargetMol. This collection encompassed 2,040 approved drugs, 5,370 bioactive compounds, and 1,680 natural compounds. Following the deduplication process, the final count of unique compounds in the library was reduced to 5,600, as detailed in **Supplementary Data 3**.

### ZINC-Cayman library for screening Aβ42 inhibitors

We conducted a screening of Aβ42 inhibitors using the ZINC-Cayman chemical library. From this collection, we selected a subset of 8,279 commercially available compounds. A total of 167 compounds were excluded due to incompatibility with RDKit loading processes. For the remaining compounds, we computed several molecular properties using RDKit and other tools to assess drug-likeness and complexity: (i) Quantitative Estimation of Drug-likeness (QED), which was calculated with RDKit; the QED score is a composite metric based on molecular descriptors such as molecular weight (MW=0.66), octanol-water partition coefficient (ALOGP=0.46), number of hydrogen bond acceptors (HBA=0.05), number of hydrogen bond donors (HBD=0.61), polar surface area (PSA=0.06), number of rotatable bonds (ROTB=0.65), number of aromatic rings (AROM=0.48), and structural alerts for reactivity (ALERTS=0.95). (ii) CNS MPO Score (cns_guacamol), which was calculated using GuacaMol; The CNS MPO score is indicative of the suitability of a compound for central nervous system activity. (iii) Complexity Score (complexity_rdkit), which was calculated with RDKit; This score evaluates the structural complexity of the molecules.

### Screening strategy of EGFR^LTC^ inhibitors

When selecting candidate ligands from the TargetMol library for binding to EGFR^LTC^, we first considered the predicted binding activity. We normalized the predicted affinities from 11 models, including the base model (Model B) and fine-tuned models (Model FT1 to FT10), and defined an overall affinity score as the minimal predicted affinity (normalized) among the 11 models (see Supplementary Information). In screening the candidate ligands, we also took into account the binding location of the ligands. We calculate the predicted distances of the ligand to residues K745, E762, D855, the three functional residues of EGFR, and normalized these distances to obtain a distance score. We screened molecules based on the following criteria: overall affinity score > 0.3 (i.e., top 50 results) or affinity score of any model rank < 10; and all distance scores are less than 0.5, to ensure that the binding location of the small molecule is not too far from the target region. Following these criteria, we obtained 12 candidates (**Extended Data Table 3**).

### Reweighting state populations of the ABL kinase

ABL exists in three different states, with state A corresponding to PDB 6XR6, state I_1_ corresponding to PDB 6XR7, and state I_2_ corresponding to PDB 6XRG. Each PDB file contains a set of 𝑁_conf_ = 20 conformations. We use the average residue distance among the 20 conformations in each state as the distance matrix 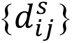. Specifically, 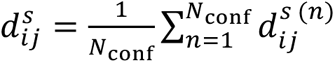, where state 𝑠 ∈ {A, I_1_, I_2=_}, and 𝑖, 𝑗 ∈ {1, … , 𝑁_res_ = 287}. The conformational ensemble of the ABL kinase can be simplified and represented by the weights 𝐰 = [𝑤^A^, 𝑤^I_1_^, 𝑤^I_2_^] of the three states. Consequently, the distance matrix 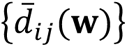 of the conformational ensemble can be expressed as a weighted average: 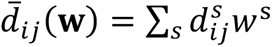. During the reweighting process, we optimize the weights 𝐰 to minimize the mean squared error (MSE) between the protein distance matrix 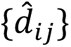 predicted by Ligand-Transformer and the distance matrix 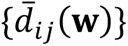 of the conformational ensemble. The optimized weights 𝐰^∗^are considered to represent the conformational population corresponding to the Ligand-Transformer predicted distance matrix. Specifically, we use Sequential Least Squares Programming (SLSQP) from SciPy to solve the following optimization problem with constraints and bounds:

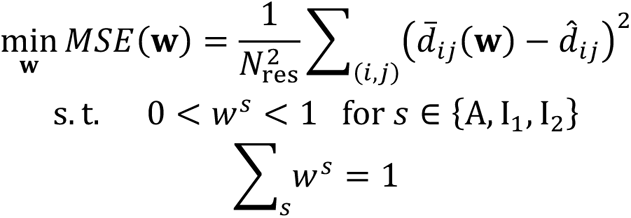

## Baselines

### HAC-Net

We obtained the code for HAC-Net from the official repository at https://github.com/gregory-kyro/HAC-Net, and used their pre-processed input HDF files for the PDBbind2020 dataset. We adjusted the dataset split to align with the division we utilized for training and testing. The training process and hyperparameters adhered to their default settings.

### TankBind

We sourced the code for TankBind from https://github.com/luwei0917/TankBind. Using their dataset construction script, we constructed a dataset split for PDBbind2020 consistent with what we used in our study. We adhered to their default training settings to retrain their model on our version of the dataset.

### MONN

We obtained the code from https://github.com/lishuya17/MONN. Their dataset pre-processing discards complexes with molecules like nucleic acids and polypeptides. Therefore, we made certain modifications to their data handling code to accommodate all the data we required. We rebuilt the dataset to align with the split we used in our paper and retrained their model using their default parameters.

### In vitro EGFR^LTC^ inhibition assays

The HTRF KINASE-TK kit assay (Cat#62TK0PEJ) was purchased from PerkinElmer. Compounds were diluted in a gradient using DMSO to a final dilution of 200× of the detection concentration, resulting in a final DMSO content of 0.5% in the assay system. 25 nL of the compound was transferred into a 384-well reaction plate (784075, Greiner) using Echo655. A kinase solution was prepared using 1× kinase reaction buffer (5× Buffer, 5 mM MgCl_2_, 1 mM DTT, 1 mM MnCl_2_) at a working concentration of 0.7 nM, and 2.5 μL of the kinase solution was transferred to the 384-well reaction plate. The plate was centrifuged at 1000 rpm for 1 min and incubated at 25 °C for 10 min. A mixture of substrate (TK substrate, working concentration: 1 μM) and ATP (working concentration: 0.2 μM) was prepared in the kinase reaction buffer, and 2.5 μL of the substrate and ATP mixture was added to the reaction plate to initiate the reaction. Then, the plate was centrifuged at 1000 rpm for 1 min, and sealed with a sealing film and incubated at 25 °C for 50 min. A mixture of 2× XL665 and antibody detection reagent was prepared in the detection buffer. 5 μL of the kinase detection reagent was added to each well of the 384-well reaction plate, which was then centrifuged at 1000 rpm for 1 min and incubated at 25 °C for 60 min. Fluorescence signals at 620 nm (Cryptate) and 665 nm (XL665) were measured using a microplate reader. Each reaction was performed in duplicate. The percentage of kinase inhibition induced by the compounds was quantified using the equation: 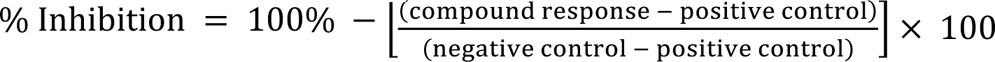. To evaluate the potency of the inhibitors, IC_50_ values and dose-response curves were generated using GraphPad Prism 7.0 software. This was achieved by fitting the calculated percentage inhibition and the logarithm of the compound concentrations to a variable slope (four-parameter) nonlinear regression model. The model is expressed by the equation: 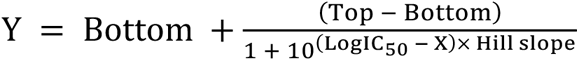. where X denotes the log of the inhibitor concentration and Y represents the percentage inhibition. Furthermore, the percentage of enzymatic activity in the presence of inhibitors, expressed as % Control, is derived by subtracting the % Inhibition from 100%.

### Aβ42 aggregation kinetics

Solutions of monomeric Aβ42 were prepared by dissolving the lyophilized Aβ42 peptide in 6 M guanidinium hydrocholoride (GuHCl). Monomeric forms were purified from potential oligomeric species and salt using a Superdex 75 10/300 GL column (GE Healthcare) at a flowrate of 0.5 mL/min, and were eluted in 20 mM sodium phosphate buffer, pH 8 supplemented with 200 µM EDTA and 0.02% NaN3. The centre of the peak was collected and the peptide concentration was determined from the absorbance of the integrated peak area using ε280 = 1490 l mol^-1^ cm^-1^. The obtained monomer was diluted with buffer to the desired concentration and supplemented with 20 μM thioflavin T (ThT) from a 2 mM stock. Each sample was then pipetted into multiple wells of a 96-well half-area, low-binding, clear bottom and PEG coated plate (Corning 3881), 80 µL per well, in the absence and the presence of 20 μM of small molecules (1% DMSO). Assays were initiated by placing the 96-well plate at 37 °C under quiescent conditions in a plate reader (Fluostar Omega, Fluostar Optima or Fluostar Galaxy, BMGLabtech, Offenburg, Germany). The ThT fluorescence was measured through the bottom of the plate using a 440 nm excitation filter and a 480 nm emission filter.

### Biolayer interferometry to measure the binding to Aβ42

10074-G5 was obtained from Cambridge Bioscience, Cayman Chemical, UK. The other small molecules were purchased from MolPort (Riga, Latvia) or Mcule (Budapest, Hungary). The small molecules were firstly dissolved in DMSO, vortexed, and then filtered using 0.02 mm filter. They were then diluted in freshly prepared buffer solutions (20 mM sodium phosphate buffer (pH 8) and 200 mM EDTA) to make final concentrations of 1, 2, 3, 4 and 5 µM with 1% DMSO. The DMSO concentration was kept at 1% in the control solutions in all experiments. Super streptavidin biosensors (Sartouris, UK) were incubated with N-terminally biotinylated Aβ42 monomers (15 μg/ml; Generon, AnaSpec, UK) by placing the tips in the monomeric Aβ42 solution at 5 °C overnight. A parallel set of biosensors were incubated in biocytin at the same conditions as a negative control. The sensors were then rinsed in buffer solution for three hours at room temperature to rinse off any potential fibrils on the surface. Octet Red 96, Fortebio BLI-based assay with a stirring speed of 1000 revolutions/min (rpm) and temperature of 27 °C were used in black 96 well throughout the project. The association and dissociation between immobilized Aβ42 and different small molecules (at concentrations 0, 1, 2, 3, 4 and 5 µM) were monitored for 120 s and 180 s, respectively. A double reference subtraction approach was applied here to correct both the buffer baseline drift and non-specific binding. The binding of buffer to both biocytin-functionalized and Aβ42-functionalized biosensor was subtracted to account for baseline drift. The binding between biocytin-functionalized biosensors and different concentrations of small molecules were subtracted to remove the non-specific binding. Data were analysed using the software Data Analysis. By globally fitting multiple kinetic traces, the software could calculate of binding affinity constants (*K*_d_) under a 1:1 binding model. The *K*_d_ values were determined based on the ratio of the association rate constant (*k*_on_) and the dissociation rate constant (*k*_off_).

---

**Extended Data Figure 1.**
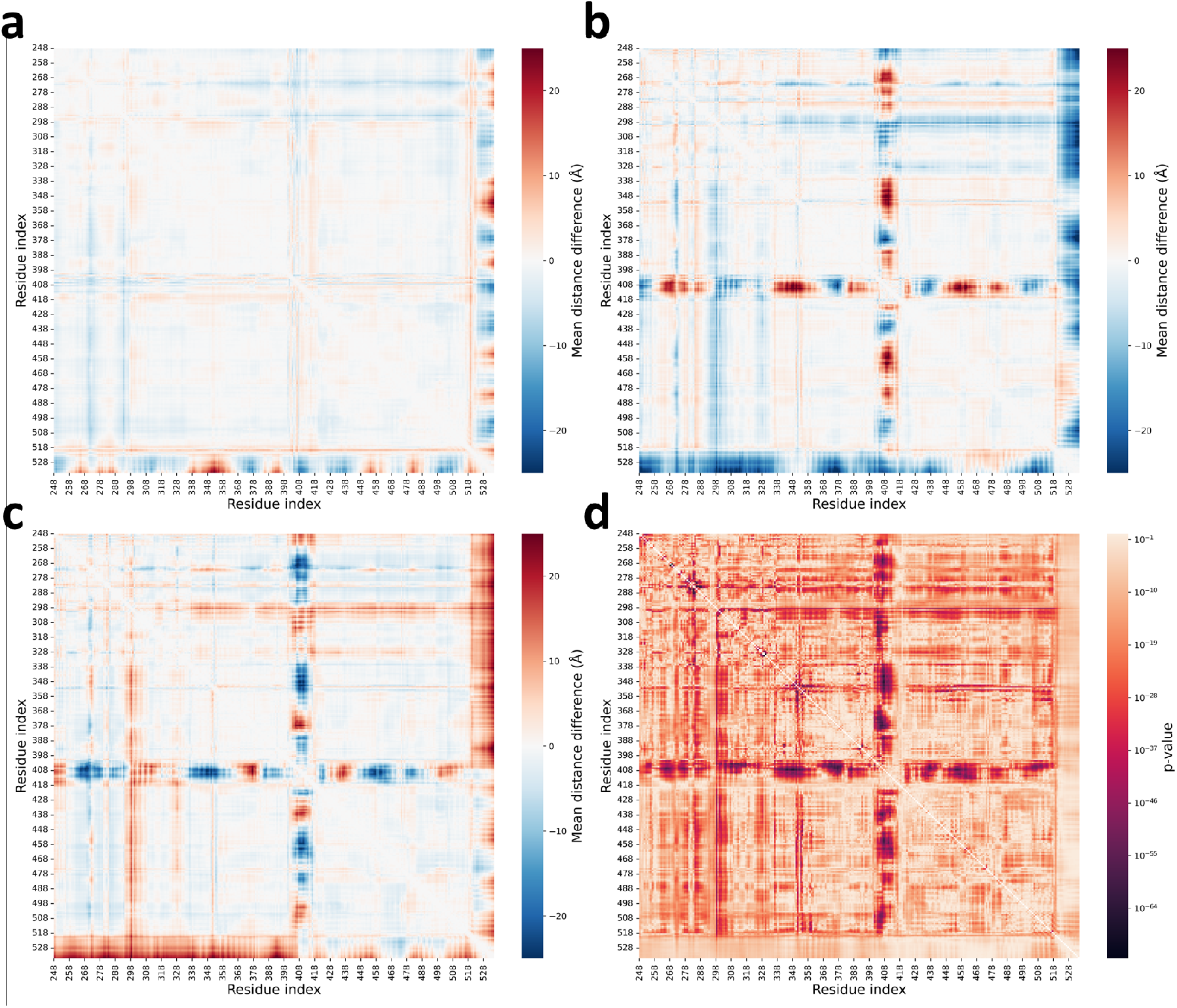
Comparative analysis of distance matrices of the states of ABL. **(a-c)** Difference distance matrices illustrating the structural variations between the three states of ABL: **(a)** Difference distance matrix of state I_1_ minus state A. **(b)** Difference distance matrix of state I_2_ minus state A. **(c)** Difference distance matrix of state I_1_ minus state I_2_. **(d)** F-statistic values for the variation in distance distribution across each pair of residues when comparing the three states. The color intensity correlates with the significance of the distance variation, with the accompanying scale indicating p-values.

**Extended Data Figure 2.**
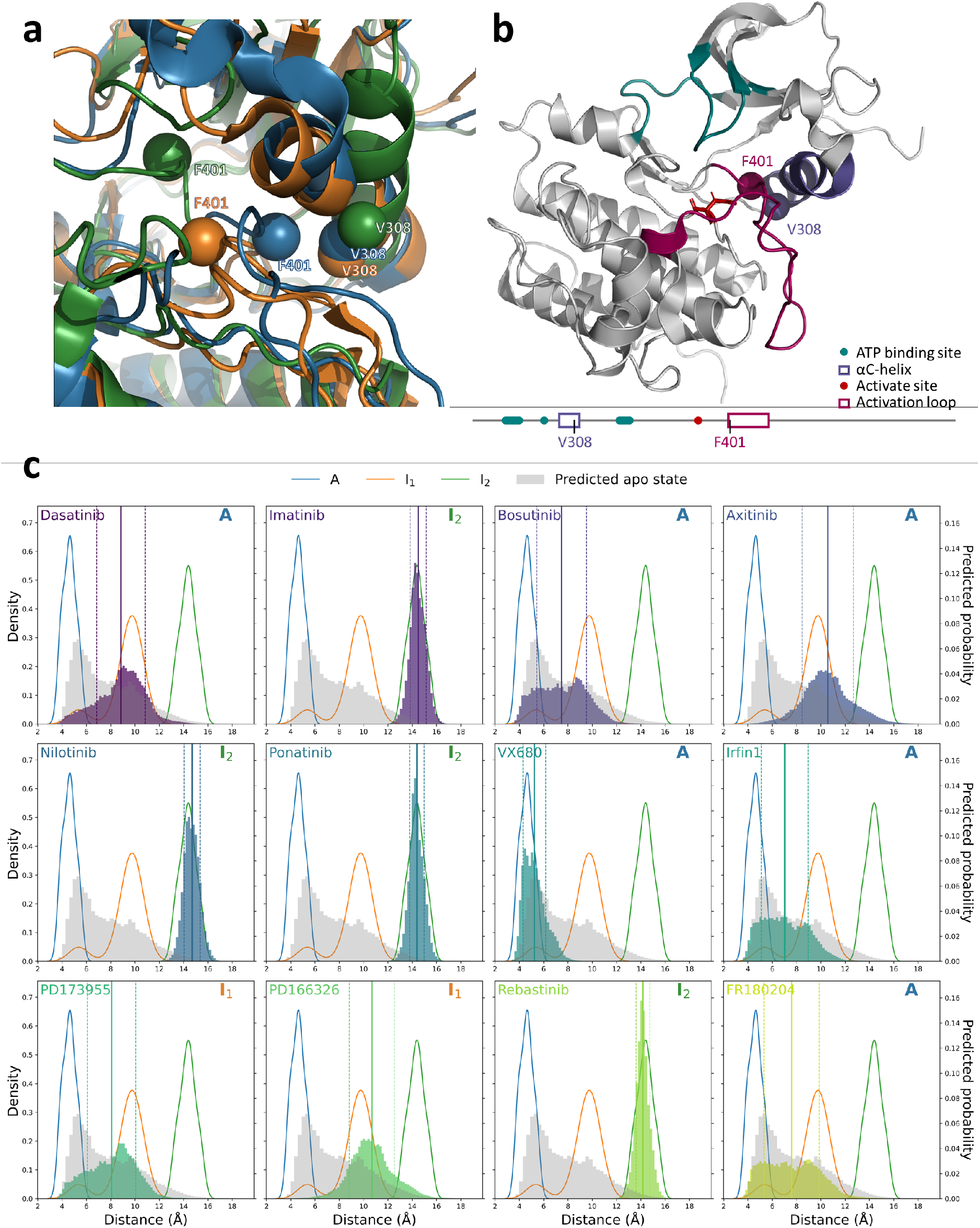
Analysis of distances between the Cβ atom of residue V308 and the Cβ atom of residue F401 of ABL. **(a)** Structural overlay highlighting the distance between residues V308 and F401 in the conformations of states A (PDB 6XR6, blue), I_1_ (PDBID: 6XR7, orange), and I_2_ (PDB 6XRG, green) of ABL. Residues are represented by spheres at the Cβ atoms, and the proteins are depicted as ribbons. **(b)** Depiction of the spatial positioning of residues V308 and F401 within the ABL kinase state A (PDB 6XR6). The sequence annotation is based on UniProt entry P00519, with the ATP binding site, αC-helix, active site, and activation loop colored in teal, violet, red, and magenta, respectively. The protein backbone is rendered as a cartoon, the active site residues as sticks, and the Cβ atoms of residues V308 and F401 as spheres. **(c)** Distance distributions between residues V308 and F401 for ABL in complex with 12 different inhibitors. Kernel density estimations (KDE) plots for the distances from 20 measured structures are shown for state A (blue), state I_1_ (orange), and state I_2_ (green). Predicted distance probability distribution of the apo state, derived from AlphaFold2 (AF2), is depicted as grey bars with a bin width of 0.3 Å. The Ligand-Transformer predicted distance probabilities for the 12 inhibitors are displayed as colored bars with a bin width of 0.19 Å. The means of predicted distances are plotted as solid lines, with dashed lines representing the standard deviation. The symbol in the upper right corner denotes which conformational state of ABL the inhibitor selectively binds to as determined by NMR analysis^23^.

**Extended Data Figure 3.**
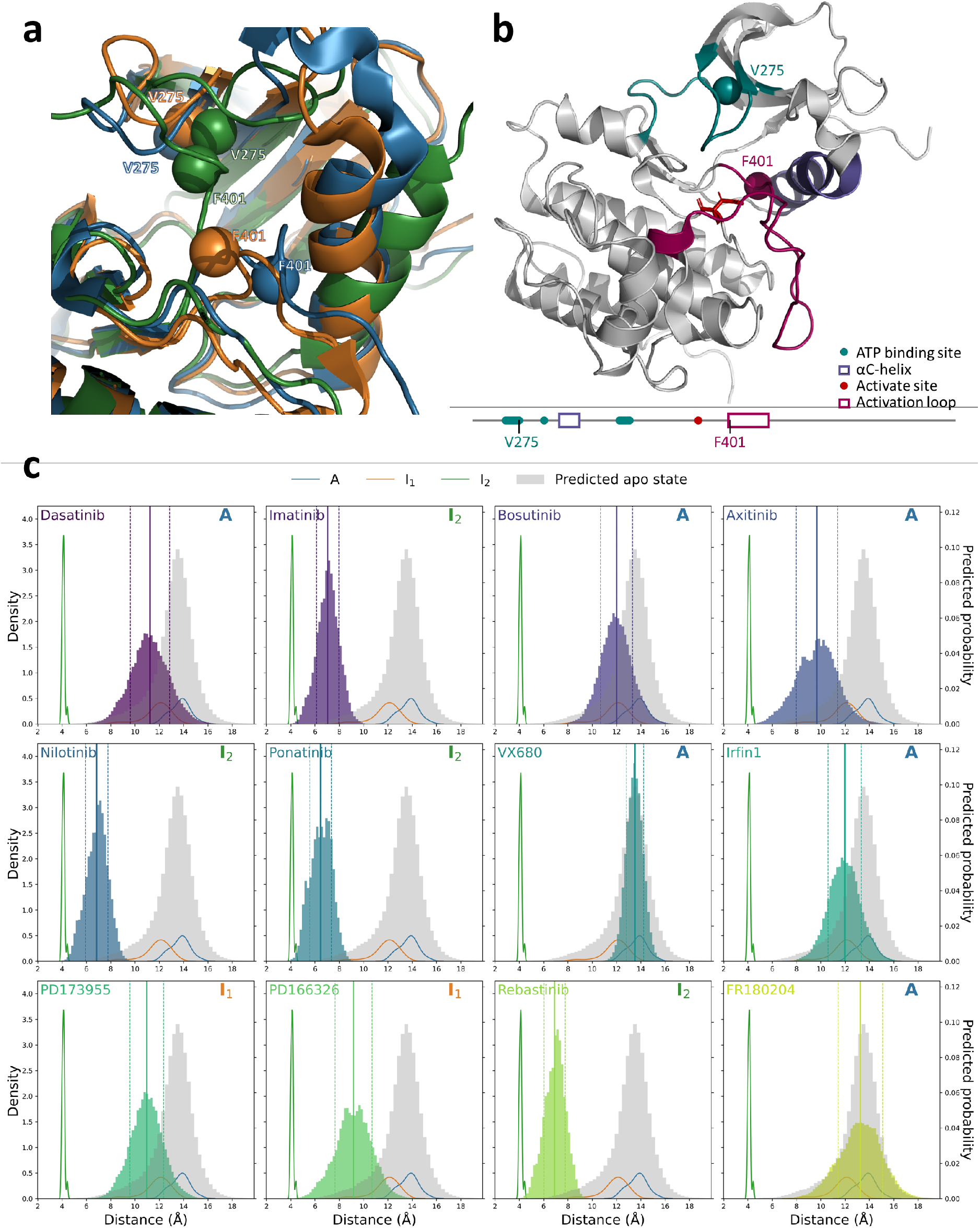
Analysis of distances between the Cβ atom of residue V275 and the Cβ atom of residue F401 of ABL kinase. **(a)** Structural overlay highlighting the distance between residues V275 and F401 in the conformations of states A (PDB 6XR6, blue), I_1_ (PDB 6XR7, orange), and I_2_ (PDB 6XRG, green) of ABL. Residues are represented by spheres at the Cβ atoms, and the proteins are depicted as ribbons. **(b)** Depiction of the spatial positioning of residues V275 and F401 within the state A of ABL (PDB 6XR6). The sequence annotation is based on UniProt entry P00519, with the ATP binding site, αC-helix, active site, and activation loop colored in teal, violet, red, and magenta, respectively. The protein backbone is rendered as a cartoon, the active site residues as sticks, and the Cβ atoms of residue V275 and F401 as spheres. **(c)** Distance distributions between residue V275 and F401 for ABL in complex with 12 different inhibitors. Kernel density estimations (KDE) plots for the distances from 20 measured structures are shown for state A (blue), state I_1_ (orange), and state I_2_ (green). Predicted distance probability distribution of the apo state, derived from AlphaFold2 (AF2), is depicted as grey bars with a bin width of 0.3 Å. The Ligand-Transformer predicted distance probabilities for the 12 inhibitors are displayed as colored bars with a bin width of 0.19 Å. The means of predicted distances are plotted as solid lines, with dashed lines representing the standard deviation. The symbol in the upper right corner denotes which conformational state of ABL the inhibitor selectively binds to as determined by NMR analysis^23^.

**Extended Data Figure 4.**
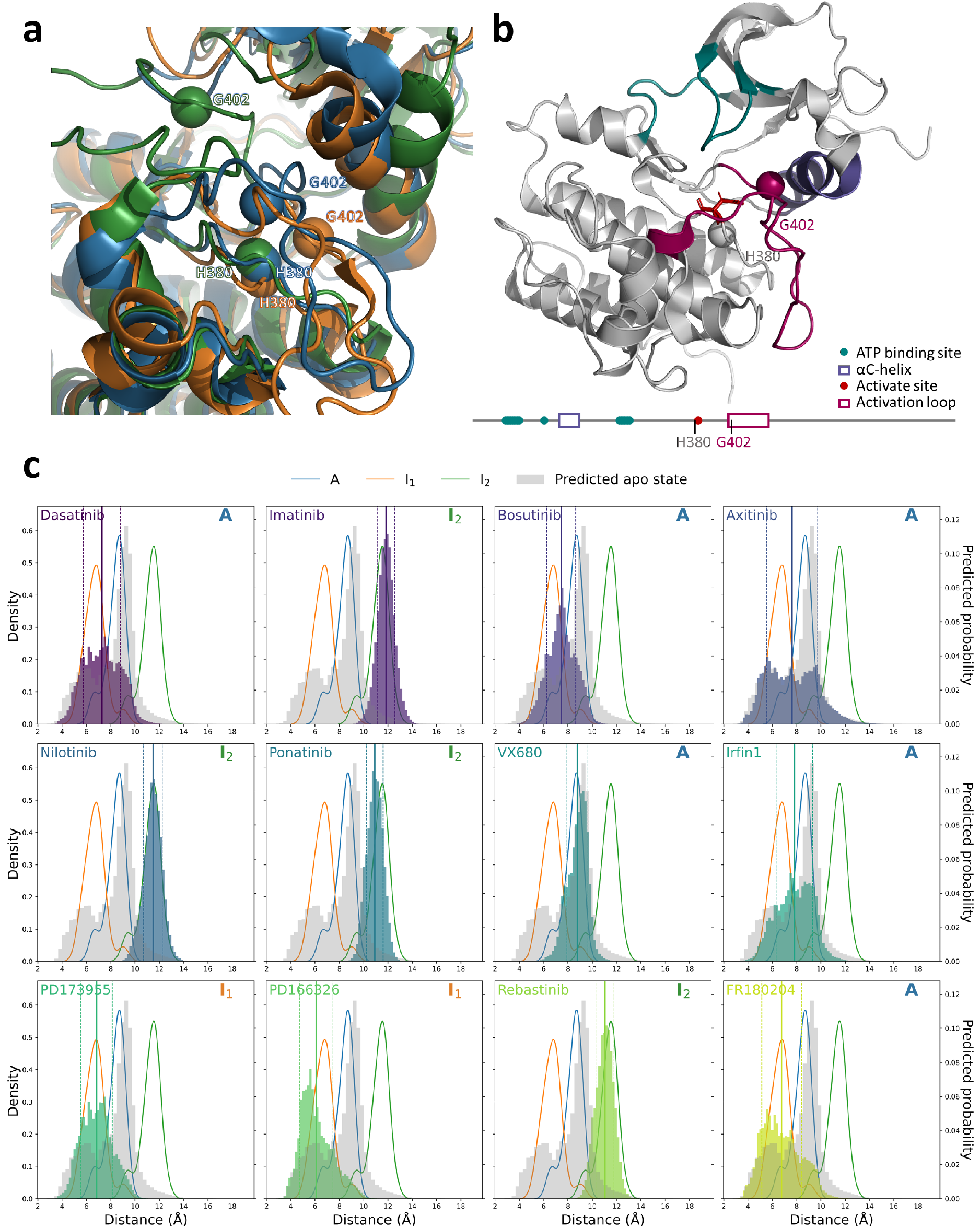
Analysis of distances between the Cβ atom of residue H380 and the Cα atom of residue G402 of ABL. **(a)** Structural overlay highlighting the distance between residues H380 and G402 in the conformations of ABL kinase states A (PDB 6XR6, blue), I_1_ (PDB 6XR7, orange), and I_2_ (PDB 6XRG, green). Residues are represented by spheres at the Cβ atoms of residue H380, Cα atoms of residue G402, and the proteins are depicted as ribbons. **(b)** Depiction of the spatial positioning of residues H380 and G402 within state A of ABL (PDB 6XR6). The sequence annotation is based on UniProt entry P00519, with the ATP binding site, αC-helix, active site, and activation loop colored in teal, violet, red, and magenta, respectively. The protein backbone is rendered as a cartoon, the active site residues as sticks, and the Cβ atom of residue V308 and the Cα atom of residue G402 as spheres. **(c)** Distance distributions between residue H380 and G402 for ABL in complex with 12 different inhibitors. Kernel density estimations (KDE) plots for the distances from 20 measured structures are shown for state A (PDB 6XR6, blue), state I_1_ (orange), and state I_2_ (green). Predicted distance probability distribution of the apo state, derived from AlphaFold2 (AF2), is depicted as grey bars with a bin width of 0.3 Å. The Ligand-Transformer predicted distance probabilities for the 12 inhibitors are displayed as colored bars with a bin width of 0.19 Å. The means of predicted distances are plotted as solid lines, with dashed lines representing the standard deviation. The symbol in the upper right corner denotes which conformational state of ABL the inhibitor selectively binds to as determined by NMR analysis^23^.

**Extended Data Figure 5.**
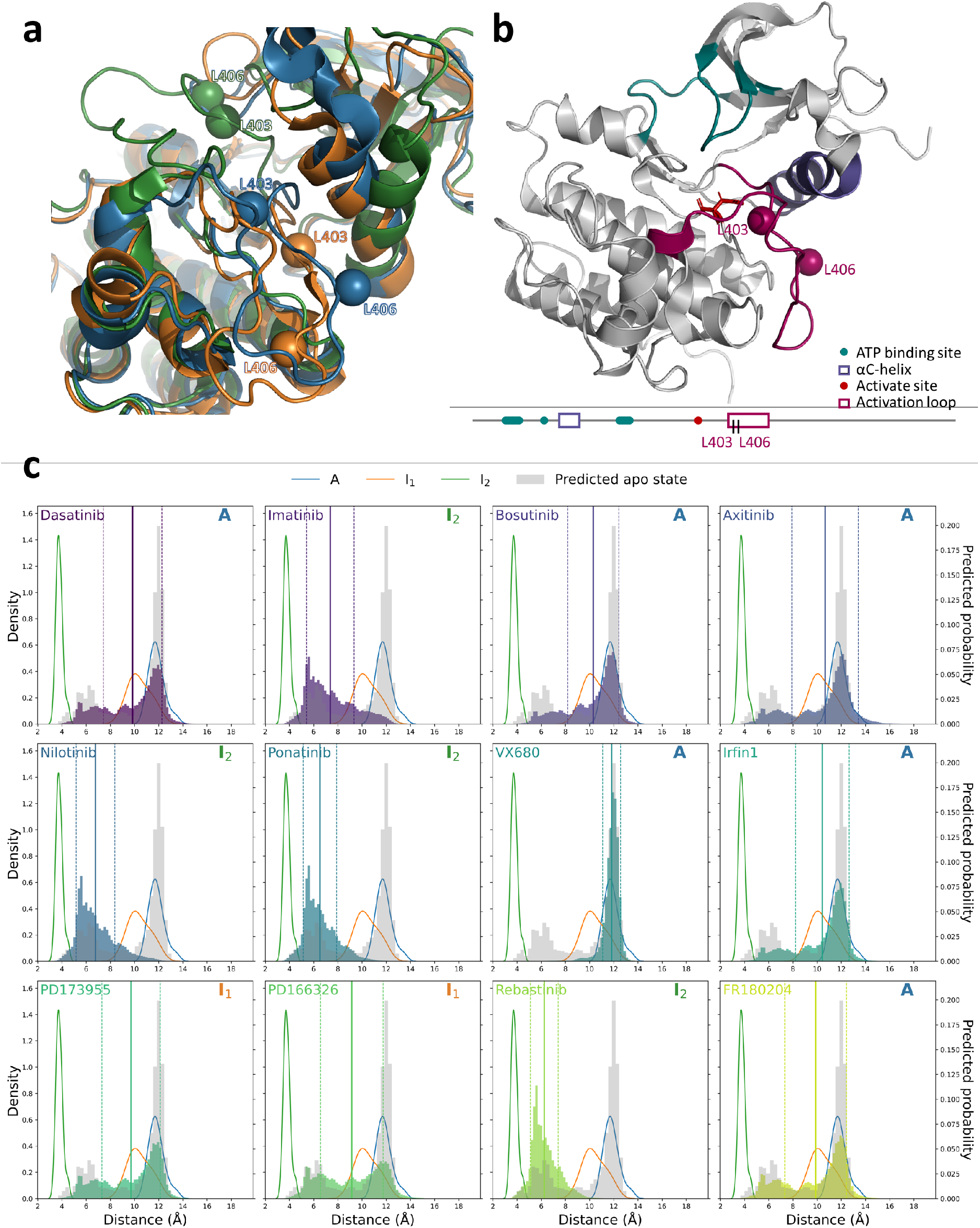
Analysis of distances between the Cβ atom of residue L403 and the Cβ atom of residue L406 of ABL. **(a)** Structural overlay highlighting the distance between residues L403 and L406 in the conformations of the states A (PDB 6XR6, blue), I_1_ (PDB 6XR7, orange), and I_2_ (PDB 6XRG, green) of ABL. Residues are represented by spheres at the Cβ atoms, and the proteins are depicted as ribbons. **(b)** Depiction of the spatial positioning of residues L403 and L406 within state A of ABL (PDB 6XR6). The sequence annotation is based on UniProt entry P00519, with the ATP binding site, αC-helix, active site, and activation loop colored in teal, violet, red, and magenta, respectively. The protein backbone is rendered as a cartoon, the active site residues as sticks, and the Cβ atoms of residue L403 and L406 as spheres. **(c)** Distance distributions between residue L403 and L406 for ABL in complex with 12 different inhibitors. Kernel density estimations (KDE) plots for the distances from 20 measured structures are shown for state A (PDB 6XR6, blue), state I_1_ (orange), and state I_2_ (green). Predicted distance probability distribution of the apo state, derived from AlphaFold2 (AF2), is depicted as grey bars with a bin width of 0.3 Å. The Ligand-Transformer predicted distance probabilities for the 12 inhibitors are displayed as colored bars with a bin width of 0.19 Å. The means of predicted distances are plotted as solid lines, with dashed lines representing the standard deviation. The symbol in the upper right corner denotes which conformational state of ABL the inhibitor selectively binds to as determined by NMR analysis^23^.

**Extended Data Figure 6.**
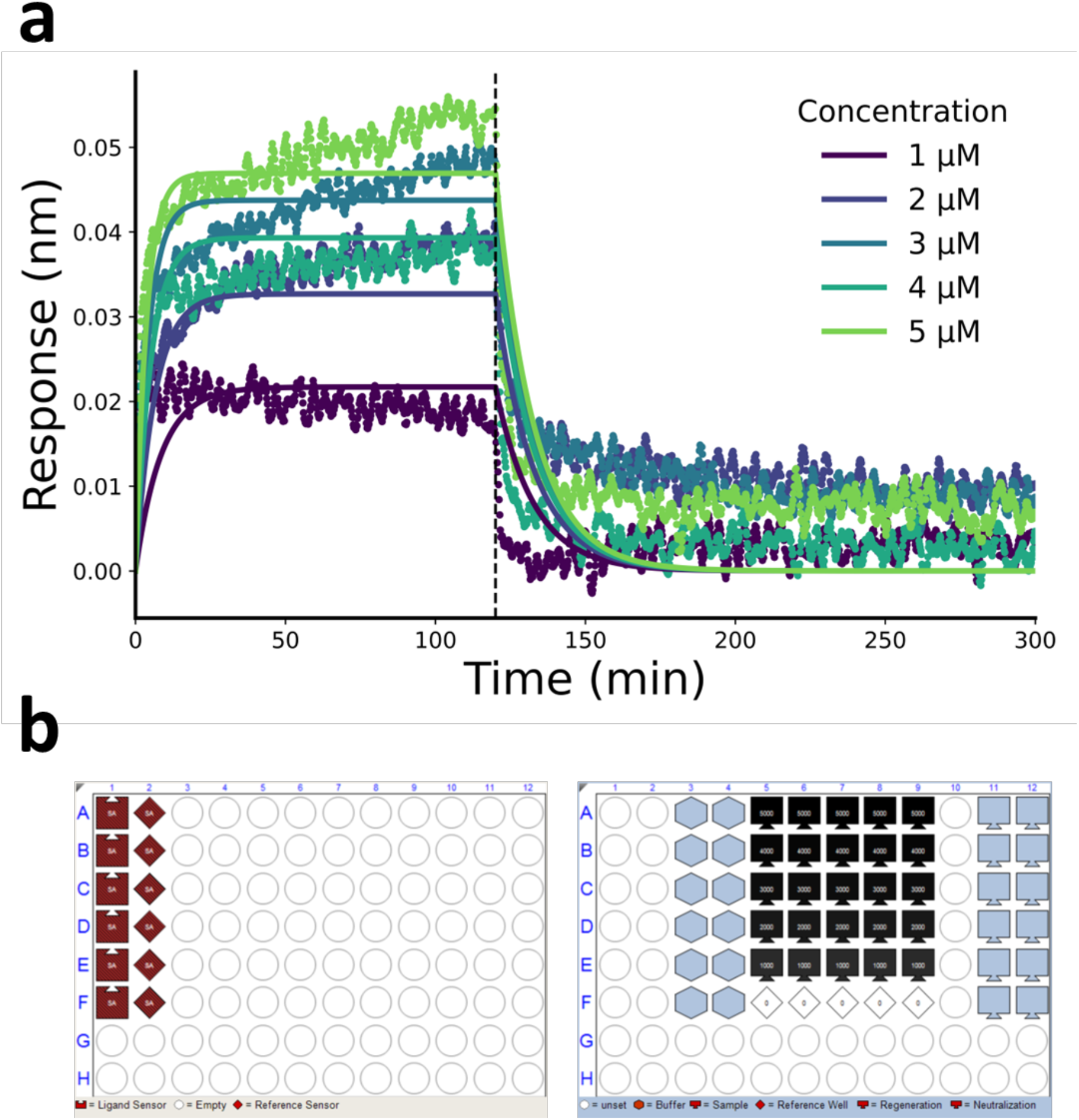
BLI measurements setup and the result for compound 10074-G5. (a) BLI measurements showing the binding kinetics of compound 10074-G5 to Aβ42-functionalized surfaces at different concentrations. The baseline drift was corrected, and global fitting to one-phase association and dissociation models provided the kinetic rates. The *k*_on_ was determined to be 3.83 × 10^4^ M^−1^s^−1^ and *k*_off_ is 7.85 × 10^-2^ M^−1^s^−1^, which gave a *K*_d_ value of 2.05 ± 0.15 μM. (b) Sensor tray and sample data. In the sensor tray, A1, B1, C1, D1, E1and F1 are the sample sensors incubated with Aβ42; A2, B2, C2, D2, E2 and F2 are the control sensors incubated with biocytin. In the sample data, the sensors were placed in 10074-G5, D4, D9, D3, and D9, respectively at five different concentrations.

**Extended Data Table 1.**
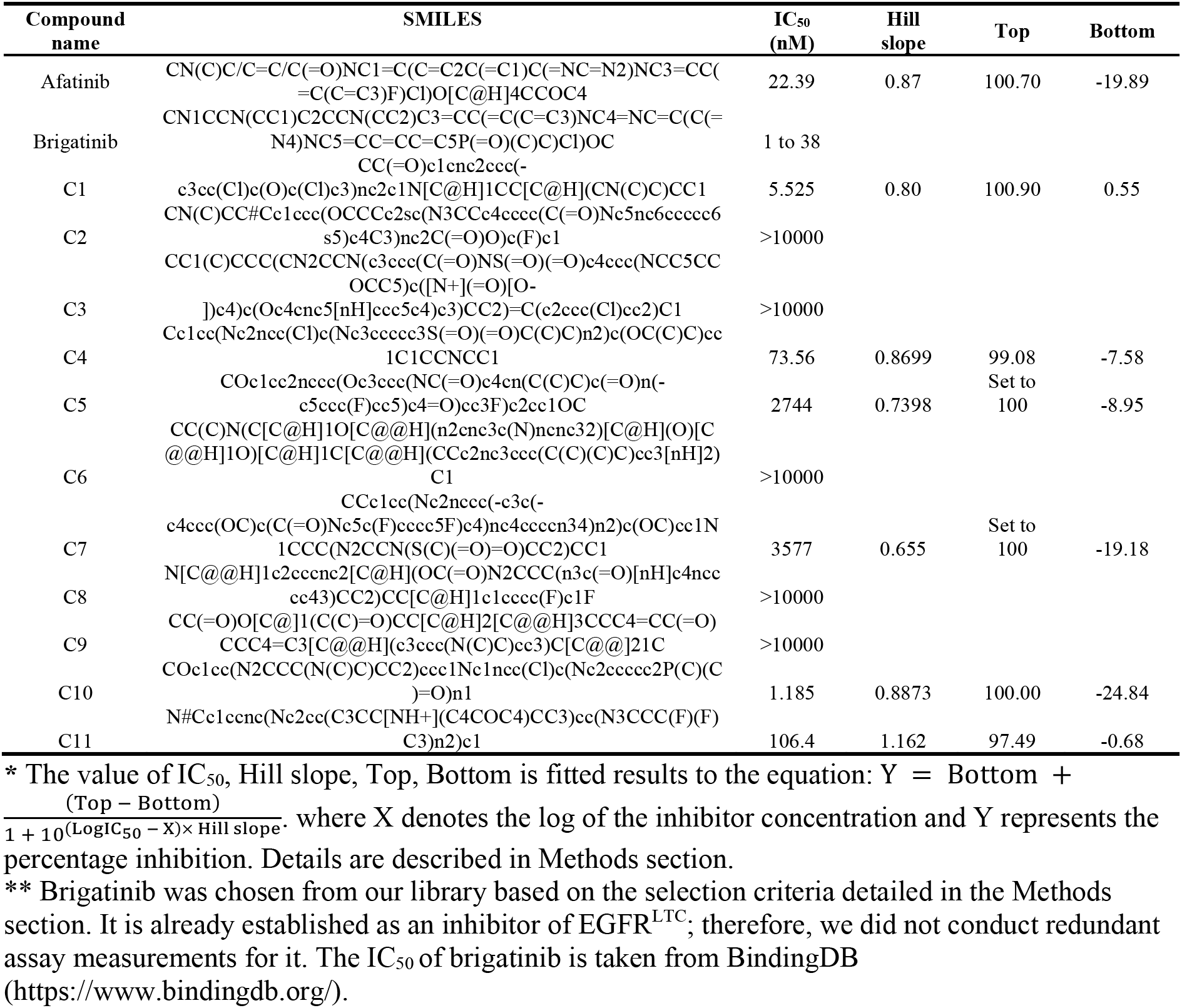
In vitro EGFR^LTC^ inhibition assays for afatinib and 12 selected ligands predicted by Ligand-Transformer.

**Extended Data Table 2.** Predicted populations of three conformational states of ABL in complex with ligands.

| Compound name | Measured state <sup>a</sup> | Predicted state <sup>b</sup> | $w^A$ <sup>c</sup> | $w^{I_1}$ <sup>c</sup> | $w^{I_2}$ <sup>c</sup> | MSE (Å) <sup>d</sup> |
| --- | --- | --- | --- | --- | --- | --- |
| Dasatinib | A | A | 0.569 | 0.359 | 0.072 | 0.313 |
| Imatinib | I<sub>2</sub> | I<sub>2</sub> | 0.254 | 0.259 | 0.487 | 0.406 |
| Bosutinib | A | A | 0.478 | 0.434 | 0.088 | 0.329 |
| Axitinib | A | I<sub>1</sub> | 0.394 | 0.41 | 0.196 | 0.340 |
| Nilotinib | I<sub>2</sub> | I<sub>2</sub> | 0.263 | 0.24 | 0.497 | 0.387 |
| Ponatinib | I<sub>2</sub> | I<sub>2</sub> | 0.273 | 0.24 | 0.486 | 0.394 |
| VX680 | A | A | 0.655 | 0.284 | 0.061 | 0.317 |
| Irfin1 | A | A | 0.607 | 0.316 | 0.076 | 0.317 |
| PD173955 | I<sub>1</sub> | I<sub>1</sub> | 0.44 | 0.468 | 0.091 | 0.320 |
| PD166326 | I<sub>1</sub> | I<sub>1</sub> | 0.404 | 0.496 | 0.101 | 0.323 |
| Rebastinib | I<sub>2</sub> | I<sub>2</sub> | 0.227 | 0.264 | 0.509 | 0.392 |
| FR180204 | A | A | 0.552 | 0.372 | 0.076 | 0.314 |
| apo <sup>e</sup> | A | A | 0.518 | 0.303 | 0.179 | 0.299 |

<sup>a</sup> Conformational state of ABL [A, I<sub>1</sub>, I<sub>2</sub>] the inhibitor selectively binds to as determined by NMR analysis<sup>23</sup>.
<sup>b</sup> Predominant conformational state of ABL inferred to the predicted population.
<sup>c</sup> Predicted population of specific conformational state of ABL according to the reweighting process detailed in the Methods section.
<sup>d</sup> Mean squared error (MSE) between the protein distance matrix $\{\hat{d}_{ij}\}$ predicted by Ligand-Transformer and the distance matrix $\{\bar{d}_{ij}(\mathbf{w})\}$ of the reweighted conformational ensemble. Details are described in the Methods section.
<sup>e</sup> Apo state of ABL, using distance matrix predicted by AF2.

**Extended Data Table 3.**
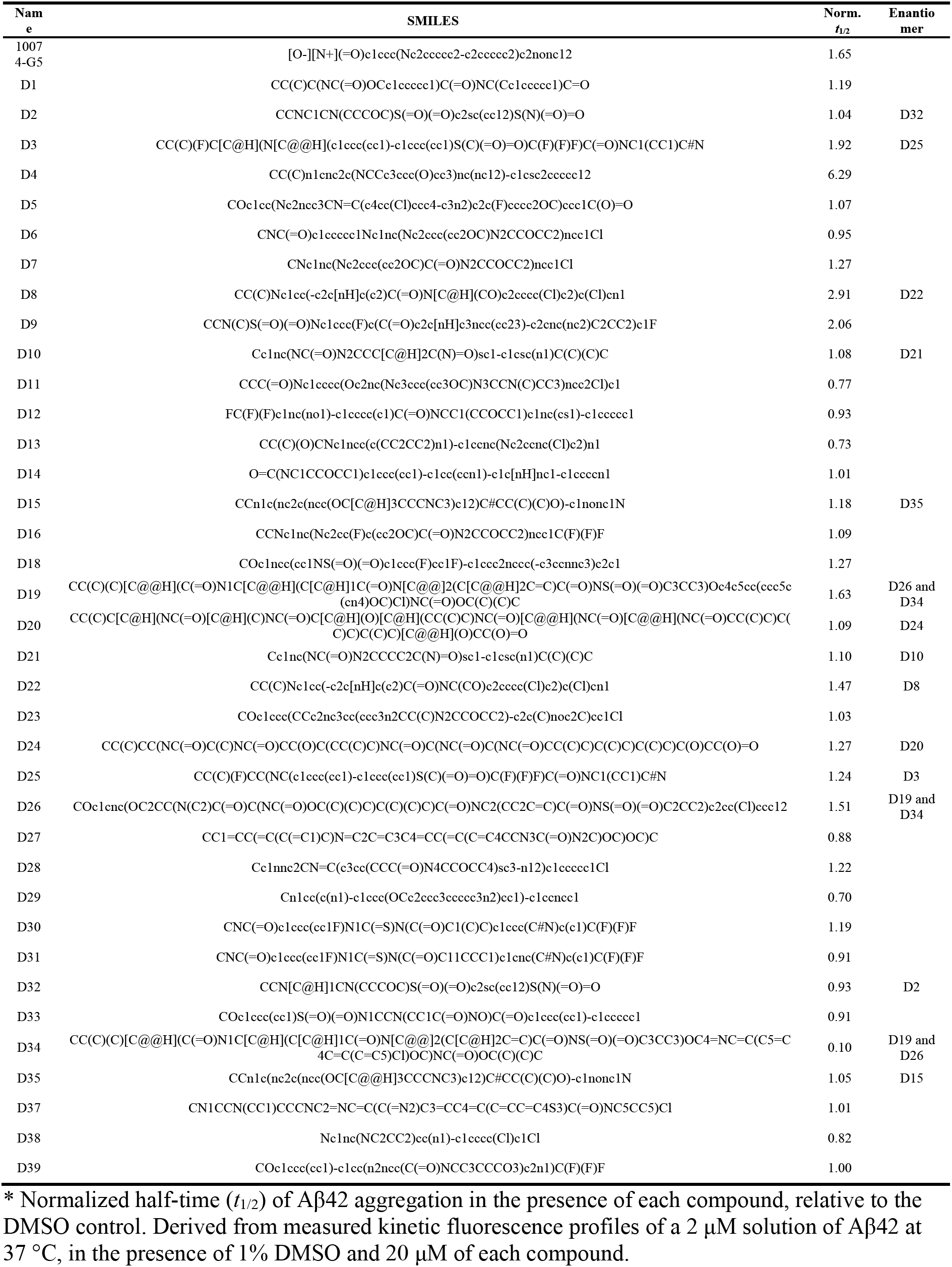
Effect on Aβ42 aggregation of 39 tested compounds.

**Supplementary Data 1. PDBbind2020-subset dataset.** This dataset comprises the training, validation, and test set split from PDBbind_10k. It includes columns for PDB ID, measured binding affinity in pKd units, protein sequences, and SMILES representations for the ligands.

**Supplementary Data 2. EGFR^LTC^-290 dataset.** This dataset consists of inhibitors targeting EGFR^LTC^, compiled from various literature sources. It provides SMILES representations for each inhibitor, alongside their measured IC_50_ values in nanomolar units. An additional "allosteric" column categorizes the inhibitor type, with ’0’, ’1’, and ’0+1’ denoting orthosteric, allosteric, and dual inhibitors, respectively.

**Supplementary Data 3. TargetMol library for screening EGFR^LTC^ inhibitors.** This dataset lists the in-stock TargetMol library utilized for screening potential EGFR^LTC^ inhibitors. It includes SMILES representations of the compounds and their original categories within TargetMol. Additionally, the dataset provides information on the maximum Tanimoto similarity for each compound compared to those in the EGFR^LTC^-290 dataset.

**Supplementary Data 4. ZINC-Cayman library for screening Aβ42 inhibitors.** This dataset lists the compounds used for the screening of Aβ42 inhibitors. It details each compound name, SMILES representation, QED, CNS MPO score, and molecular complexity. The definition and calculation of these properties are described in the Methods section.

## References

1 Jumper, J. et al. Highly accurate protein structure prediction with AlphaFold. Nature 596, 583–589 (2021).

2 Evans, R. et al. Protein complex prediction with AlphaFold-multimer. bioRxiv, 2021.2010.2004.463034 (2021).

3 Krishna, R. et al. Generalized biomolecular modeling and design with RoseTTAFold all-atom. bioRxiv, 2023.2010.2009.561603 (2023).

4 Lin, Z. et al. Evolutionary-scale prediction of atomic-level protein structure with a language model. Science 379, 1123–1130 (2023).

5 Guo, H.-B. et al. AlphaFold2 models indicate that protein sequence determines both structure and dynamics. Sci. Rep. 12, 10696 (2022).

6 Heo, L. & Feig, M. Multi-state modeling of G-protein coupled receptors at experimental accuracy. Proteins 90, 1873–1885 (2022).

7 Robertson, A. J., Courtney, J. M., Shen, Y., Ying, J. & Bax, A. Concordance of X-ray and AlphaFold2 models of SARS-CoV-2 main protease with residual dipolar couplings measured in solution. J. Am. Chem. Soc. 143, 19306–19310 (2021).

8 Del Alamo, D., Sala, D., Mchaourab, H. S. & Meiler, J. Sampling alternative conformational states of transporters and receptors with AlphaFold2. eLife 11, e75751 (2022).

9 Stein, R. A. & Mchaourab, H. S. SPEACH_AF: Sampling protein ensembles and conformational heterogeneity with alphafold2. *PLoS Comp*. Biol. 18, e1010483 (2022).

10 Vani, B. P., Aranganathan, A., Wang, D. & Tiwary, P. AlphaFold2-rave: From sequence to Boltzmann ranking. J. Chem. Theory Comput. (2023).

11 Brotzakis, Z. F., Zhang, S. & Vendruscolo, M. AlphaFold prediction of structural ensembles of disordered proteins. bioRxiv, 2023.2001.2019.524720 (2023).

12 Liu, S., et al. Pre-training molecular graph representation with 3D geometry. *arXiv* 2110.07728 (2021).

13 Lu, W. et al. Tankbind: Trigonometry-aware neural networks for drug-protein binding structure prediction. Adv. Neural Inf. Process. Syst. 35, 7236–7249 (2022).

14 Li, S. et al. Monn: A multi-objective neural network for predicting compound-protein interactions and affinities. Cell Syst. 10, 308–322. e311 (2020).

15 Kyro, G. W., Brent, R. I. & Batista, V. S. Hac-Net: A hybrid attention-based convolutional neural network for highly accurate protein–ligand binding affinity prediction. J. Chem. Inf. Model. 63, 1947–1960 (2023).

16 Hynes, N. E. & Lane, H. A. ERBB receptors and cancer: The complexity of targeted inhibitors. Nat. Rev. Cancer 5, 341–354 (2005).

17 Lu, X. et al. Targeting EGFRL858R/T790M and EGFRL858R/T790M/c797s resistance mutations in NSCLC: Current developments in medicinal chemistry. Med. Res. Rev. 38, 1550–1581 (2018).

18 Jura, N. et al. Catalytic control in the EGF receptor and its connection to general kinase regulatory mechanisms. Mol. Cell 42, 9–22 (2011).

19 Zhang, X., Gureasko, J., Shen, K., Cole, P. A. & Kuriyan, J. An allosteric mechanism for activation of the kinase domain of epidermal growth factor receptor. Cell 125, 1137–1149 (2006).

20 Liu, Z. et al. PDB-wide collection of binding data: Current status of the PDBbind database. Bioinformatics 31, 405–412 (2015).

21 Landrum, G. RDKit: Open-source cheminformatics software. (2016).

22 Taylor, S. S. & Kornev, A. P. Protein kinases: Evolution of dynamic regulatory proteins. Trends Biochem. Sci. 36, 65–77 (2011).

23 Xie, T., Saleh, T., Rossi, P. & Kalodimos, C. G. Conformational states dynamically populated by a kinase determine its function. Science 370, eabc2754 (2020).

24 Attwood, M. M., Fabbro, D., Sokolov, A. V., Knapp, S. & Schiöth, H. B. Trends in kinase drug discovery: Targets, indications and inhibitor design. Nat. Rev. Drug Discov. 20, 839–861 (2021).

25 Witte, O. N., Dasgupta, A. & Baltimore, D. Abelson murine leukaemia virus protein is phosphorylated in vitro to form phosphotyrosine. Nature 283, 826–831 (1980).

26 Greuber, E. K., Smith-Pearson, P., Wang, J. & Pendergast, A. M. Role of ABL family kinases in cancer: From leukaemia to solid tumours. Nat. Rev. Cancer 13, 559–571 (2013).

27 Knowles, T. P., Vendruscolo, M. & Dobson, C. M. The amyloid state and its association with protein misfolding diseases. Nat. Rev. Mol. Cell Biol. 15, 384–396 (2014).

28 Hampel, H. et al. The amyloid-β pathway in Alzheimer’s disease. Mol. Psychiatry 26, 5481–5503 (2021).

29 Heller, G. T. et al. Small-molecule sequestration of amyloid-β as a drug discovery strategy for Alzheimer’s disease. Sci. Adv. 6, eabb5924 (2020).

30 Bulawa, C. E. et al. Tafamidis, a potent and selective transthyretin kinetic stabilizer that inhibits the amyloid cascade. Proc. Natl. Acad. Sci. USA 109, 9629–9634 (2012).

31 Heller, G. T., Bonomi, M. & Vendruscolo, M. Structural ensemble modulation upon small-molecule binding to disordered proteins. J. Mol. Biol. 430, 2288–2292 (2018).

32 Robustelli, P. et al. Molecular basis of small-molecule binding to α-synuclein. J. Am. Chem. Soc. 144, 2501–2510 (2022).

33 Yu, C. et al. Structure-based inhibitor design for the intrinsically disordered protein c-Myc. Sci. Rep. 6, 22298 (2016).

34 Ruan, H. et al. Computational strategy for intrinsically disordered protein ligand design leads to the discovery of p53 transactivation domain i binding compounds that activate the p53 pathway. Chem. Sci. 12, 3004–3016 (2021).

35 Follis, A. V., Hammoudeh, D. I., Wang, H., Prochownik, E. V. & Metallo, S. J. Structural rationale for the coupled binding and unfolding of the c-Myc oncoprotein by small molecules. Chem. Biol. 15, 1149–1155 (2008).

36 Lindorff-Larsen, K., Best, R. B., DePristo, M. A., Dobson, C. M. & Vendruscolo, M. Simultaneous determination of protein structure and dynamics. Nature 433, 128–132 (2005).

37 Boehr, D. D., Nussinov, R. & Wright, P. E. The role of dynamic conformational ensembles in biomolecular recognition. Nat. Chem. Biol. 5, 789–796 (2009).

38 Mittermaier, A. & Kay, L. E. New tools provide new insights in NMR studies of protein dynamics. science 312, 224–228 (2006).

39 Lane, T. J. Protein structure prediction has reached the single-structure frontier. Nat. Methods 20, 170–173 (2023).

40 Gong, G. Q. et al. A small-molecule pi3kα activator for cardioprotection and neuroregeneration. Nature, 1–10 (2023).

41 Kalliokoski, T., Kramer, C., Vulpetti, A. & Gedeck, P. Comparability of mixed IC50 data - a statistical analysis. PLoS One 8, e61007 (2013).

42 Mendez, D. et al. ChEMBL: Towards direct deposition of bioassay data. Nucleic Acids Res. 47, D930–D940 (2019).

43 Karnik, K. S., Sarkate, A. P., Tiwari, S. V., Azad, R. & Wakte, P. S. Design, synthesis, biological evaluation and in silico studies of EGFR inhibitors based on 4-oxo-chromane scaffold targeting resistance in non-small cell lung cancer (NSCLC). Med. Chem. Res. 31, 1500–1516 (2022).

