## Supplementary Information for "Sequence-based drug design using transformers"

### Ligand-Transformer framework

The overall framework of Ligand-Transformer is shown in **Figure S1**. The modelling of protein-ligand complex can be viewed as multimodal learning. In this approach, we represent the protein-ligand complex as a complete heterogeneous graph, denoted by $G = (\{\mathbf{n}_i\}, \{\mathbf{n}_k\}, \{\mathbf{e}_{ij}\}, \{\mathbf{e}_{kl}\}, \{\mathbf{e}_{ik}\})$, where $\{\mathbf{n}_i\}$ and $\{\mathbf{n}_k\}$ are the set of residues of the protein and the set of atoms of the molecule, respectively, and $\{\mathbf{e}_{ij}\}, \{\mathbf{e}_{kl}\}$ and $\{\mathbf{e}_{ik}\}$ are all the pairs of residues, all the pairs of atoms, and all the pairs of residues and atoms, respectively. The representation of protein generated by AlphaFold2[1] is fed to the protein encoding module to produce the initial graph of the protein $G^{(0)}_{\text{protein}} = \left(\left\{\mathbf{n}_i^{(0)}\right\}, \left\{\mathbf{e}_{ij}^{(0)}\right\}\right)$. Similarly, the representation of molecule generated by GraphMVP[2] is processed by ligand encoding module to generate the initial graph of the molecule $G^{(0)}_{\text{ligand}} = \left(\left\{\mathbf{n}_k^{(0)}\right\}, \left\{\mathbf{e}_{kl}^{(0)}\right\}\right)$. Then, $G^{(0)}_{\text{protein}}$ and $G^{(0)}_{\text{ligand}}$ are concatenated and padded as the initial graph of the complex $G^{(0)}$, form the basis for the subsequent processing steps of cross-modal attention module, which consists of 12 layers of transformer-like block (see the *Self-attention block* section). In each block, the node representations of in the complex graph are updated through self-attention with pair bias based on the edge representations, and the edge representations are updated based on the query-key product generated by the node representations in the self-attention module. The final node representations are fed to the affinity head, where they are pooled and used to predict the binding affinity through a fully connected regression layer. Meanwhile, the final edge representations are linearly projected to a binned distance distribution as the distance prediction. The edge set $\{\mathbf{e}_{ij}\}$ is used to predict the residue-residue distances $(\hat{d}_{ij})$, which represents the distance between Cβ atoms of residues (or Cα for glycine). The edge set $\{\mathbf{e}_{kl}\}$ is used to predict the atom-atom distances in the ligand $(\hat{d}_{kl})$, while the edge set $\{\mathbf{e}_{ik}\}$ is used to predict the residue-atom distances $(\hat{d}_{ik})$, which are represented by the distance between Cβ atoms from the residue and the atom from the ligand. The detailed information regarding the modules can be found in the Algorithm subsections, where the accompanying pseudocodes for each module are also provided.

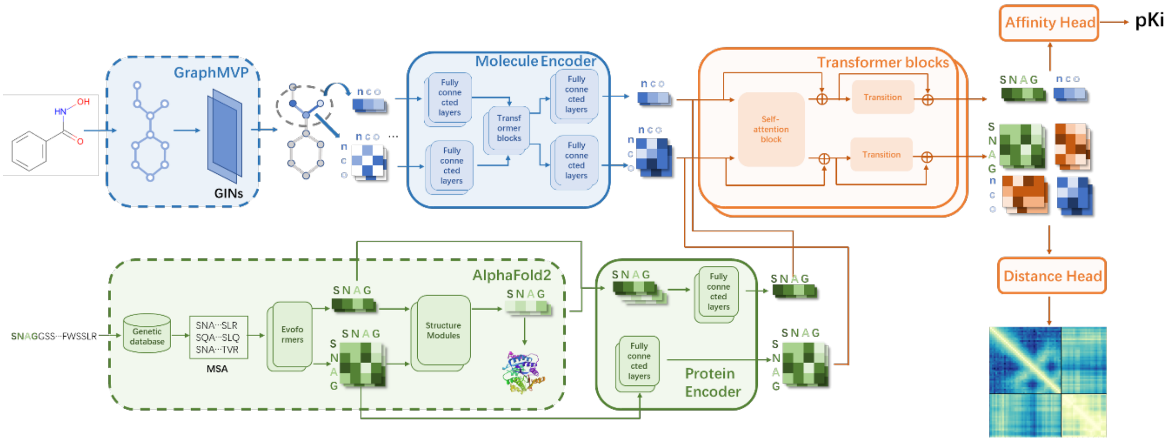


**Figure S1. Model architecture of Ligand-Transformer.** Ligand-Transformer represents a protein-ligand complex as a heterogeneous graph, incorporating residue and atom sets from the protein and ligand, respectively, along with pairwise features. The graph is formed from inputs generated by AlphaFold2 for proteins and GraphMVP for ligands, which are then re-encoded into an initial complete graph. This graph is subsequently processed through a 12-layer transformer-like network. This network updates both node and edge representations via self-attention with pair bias. The output is processed by the affinity head for binding affinity prediction and by the distance head for distance distribution prediction. Components within dotted-line boxes have fixed parameters, whereas those within solid lines are trainable.

### Benchmark test on PDBbind2020

We assessed the performance of binding affinity and distance predictions of Ligand-Transformer using the publicly available PDBbind2020[3] dataset. To work within a manageable training time, we imposed a limit on the maximum length of protein sequences to 384, and the maximum number of atoms in ligand molecules to 128. This strategy involves the selective consideration of polypeptide chains or domains situated close to the binding site (see the *Protein sequence truncation* subsection for more details). Consequently, we obtained a dataset comprised of 13,420 data points (PDBbind2020-subset, **Supplementary Data 1**). This dataset was randomly partitioned into training, validation, and testing sets, containing 10,375, 640, and 936 data points, respectively. Descriptive statistics pertaining to these partitions are presented in the *Dataset preparation* section**.**

With the training procedure described in the *Training* section, we conducted joint training for both binding affinity and distance predictions. The results of the binding affinity prediction on the test set are presented in **Figure S2a**. The majority of predictions have a low error, with the most frequent errors falling below 0.5 in p$K_d$ units (**Figure S2b**). These results indicate that the model is generally accurate in predicting the binding affinity, with a significant number of predictions closely aligning with the experimental values. For comparison, we report the results of three state-of-the-art (SOTA) models trained under identical split settings. In this test, Ligand-Transformer showed better performance across various evaluation metrics, including RMSE (root mean square error), MAE (mean absolute error), Pearson's correlation, and Spearman's correlation, with respect to the other three methods[4-6] (**Table S1**). It is worth noting that while Hac-Net[6] is a complex-based model and TankBind[4] is a pocket-based model, our model has acquired structural information solely from protein sequences.

**Table S1. Comparisons of the accuracy of binding affinity predictions with baselines.**

| Dataset | Model | Metrics | | | |
|---|---|---|---|---|---|
| | | RMSE↓ | MAE↓ | Pearson↑ | Spearman↑ |
| Validation | MONN | 1.085 | - | 0.820 | 0.818 |
| | TankBind | 1.246 | - | 0.769 | - |
| | HAC-Net | 1.240 | 0.978 | 0.756 | 0.746 |
| | Ligand-Transformer | **1.076** | **0.817** | **0.825** | **0.832** |
| Test | MONN | 1.234 | - | 0.775 | 0.776 |
| | TankBind | 1.269 | - | 0.775 | - |
| | HAC-Net | 1.265 | 0.995 | 0.761 | 0.752 |
| | Ligand-Transfomer | **1.178** | **0.861** | **0.800** | **0.803** |

The following evaluation metrics are reported: root-mean-square error (RMSE) in p$K_d$ units, mean absolute error (MAE) in p$K_d$ units, Pearson's correlation, and Spearman's correlation. The best result for each split and metric is marked in bold. All models were trained and tested on the identical split of PDBbind_10k, which was constructed in this study.

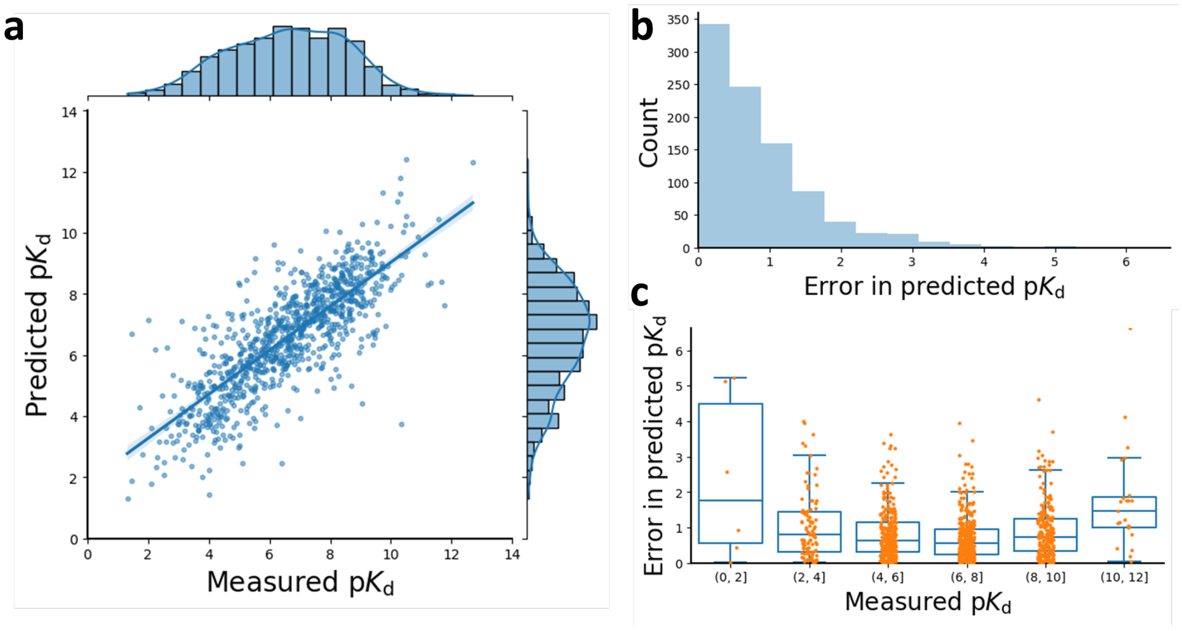


**Figure S2: Evaluation of Ligand-Transformer affinity predictions on the PDBbind_10k test set. (a)** Scatter plot with regression analysis of measured vs predicted p$K_d$ values, featuring a linear regression line with 95% confidence intervals obtained from bootstrapping. Marginal histograms and kernel density estimate plots illustrate the distributions of the measured and predicted values. **(b)** Histogram detailing the distribution of the absolute prediction errors for p$K_d$ values. **(c)** Box plots segmenting the absolute prediction errors for p$K_d$ values across varying ranges of measured p$K_d$.

**Figure S2c** provides information about the performance of the prediction model across different ranges of p$K_d$ values. The performance of the model is more robust and accurate for median p$K_d$ values (ranging from 4 to 10). The deterioration of the performance in the extreme ranges could stem from biases inherent in the PDBbind dataset, which tends to include molecules with moderate to strong binding affinities, thus under-representing ligands with weak or no binding. Additionally, compounds with very high binding affinities are infrequently discovered.

For the distance predictions, since analysing the distance predictions of all pairs is expensive, we measured the mean absolute error (MAE) of the predicted $\hat{d}_{ij}$, $\hat{d}_{kl}$, and $\hat{d}_{ik}$ of each protein-ligand complex. Around 95% of the MAEs of $\hat{d}_{ij}$ in the test results were under 0.5 Å, around 95% of the MAEs of $\hat{d}_{kl}$ were under 1 Å, and 95% of the MAEs of $\hat{d}_{ik}$ were under 2 Å (**Figure S2a**). Additionally, we found that our confidence module (see **Algorithm 8**), which predicts the mean absolute error (pMAE) of distances, is effective in estimating errors (**Figure S2c**). Specifically, the Pearson's correlation coefficients between the pMAE and true MAEs of $\hat{d}_{ij}$, $\hat{d}_{kl}$, and $\hat{d}_{ik}$ were 0.90, 0.73, and 0.56, respectively. The distance prediction errors were higher for low-resolution data (as determined by NMR spectroscopy or resolution ≥ 3 Å) compared with high-resolution data (**Figure S2d**). Low-resolution structures often have a greater degree of uncertainty in atomic positions, which naturally leads to higher errors in predicted distances within protein-ligand complexes. Such biases indicate that part of the prediction inaccuracies might originate from experimental errors or the inherent structural uncertainties. Moreover, the fact that the model can estimate its own distance prediction errors suggests that it has learned to recognize specific features within the protein-ligand complex that might be indicative of conformational variability. However, the estimation of distance prediction error does not translate into an ability to estimate binding affinity prediction error (**Figure S3d**).

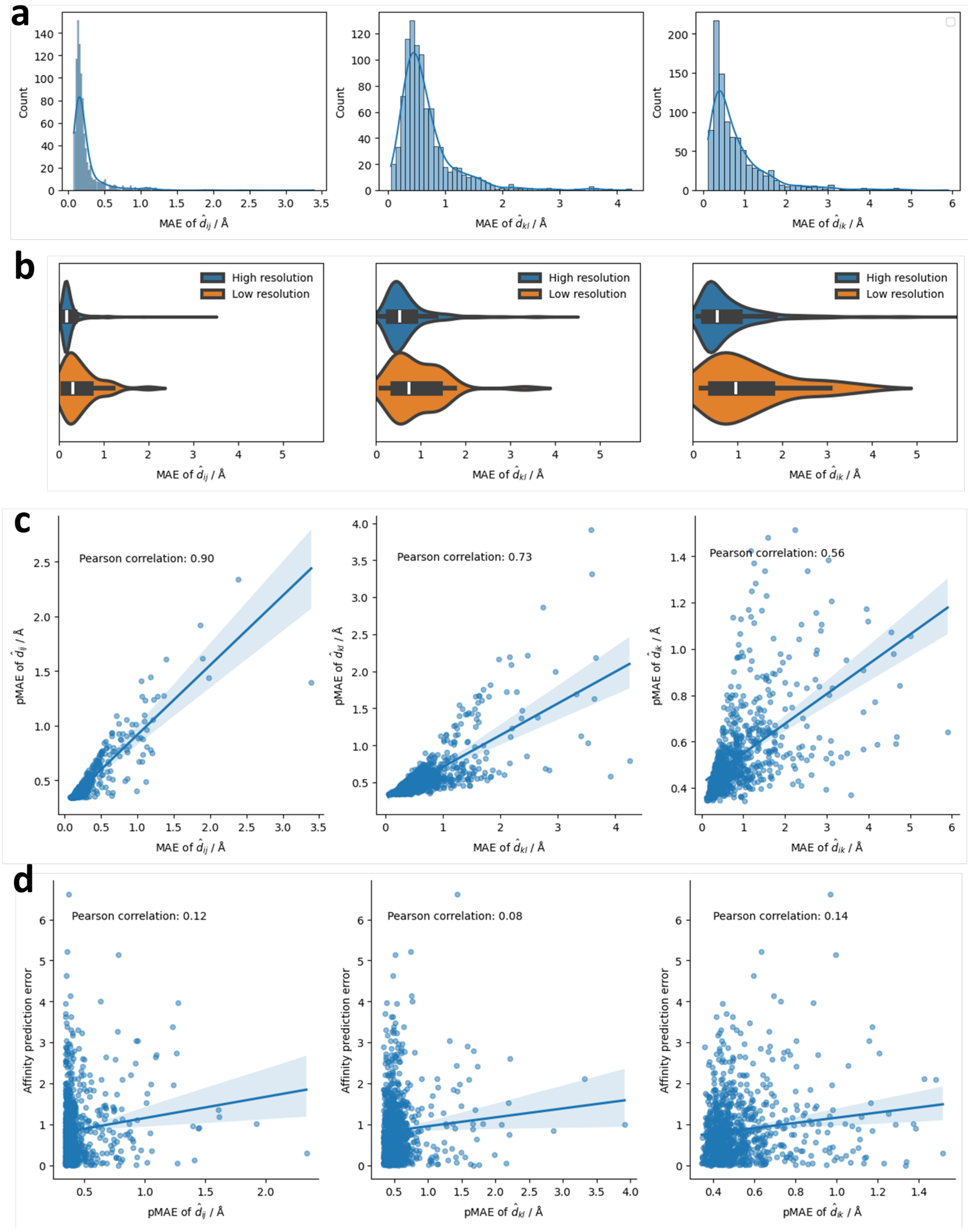


**Figure S3. Evaluation of Ligand-Transformer distance predictions on the PDBbind_10k test set. (a)** Histograms depicting the distribution of mean absolute errors (MAEs) in distance predictions by Ligand-Transformer for individual complexes within the test set. The subplots from left to right illustrate the MAEs for residue-residue distances ($\hat{d}_{ij}$, defined as the distance between the Cβ atoms of the residues, or the Cα atom in the case of glycine), ligand atom-atom distances ($\hat{d}_{kl}$), and residue-ligand atom distances ($\hat{d}_{ik}$, indicating distances between the Cβ atoms of the residues, or the Cα in the case of glycine, and ligand atoms). **(b)** Comparative analysis of distance prediction errors for structures with low resolution (either NMR-based or with resolution ⩾ 3 Å, n=39) versus high-resolution structures (resolution < 3 Å, n=897). **(c)** Scatter plots of the correlation between the predicted MAEs (pMAEs) for distances and the actual MAEs, with pMAEs calculated as the average of all predicted absolute errors (pAEs) from the confidence module (refer to **Algorithm 8** for details). The pMAEs are plotted for three distance categories: intra-protein residue-residue ($\hat{d}_{ij}$), intra-ligand atom-atom ($\hat{d}_{kl}$), and protein-ligand residue-atom ($\hat{d}_{ik}$). **(d)** Correlation plots showing the relationship between pMAEs of distance predictions and the absolute errors in pKd prediction, in log10 Molar units. The shaded areas representing 95% confidence intervals of the regression lines.

We validated the generalisation ability of the model by analysing the performance on unseen protein sequences and ligands in its training set. We found that the protein sequences in the test set have considerable overlap with the training set, and the ligand chemical space of the test set exhibits more distinct characteristics compared to the training set (**Figure S4a**,**b**). We also observed that there are no significant prediction error biases when dealing with unseen protein sequences, ligands, or combinations of both (**Figures S4c-f**). These results suggest that the model exhibits a good capability to handle new targets or ligands.

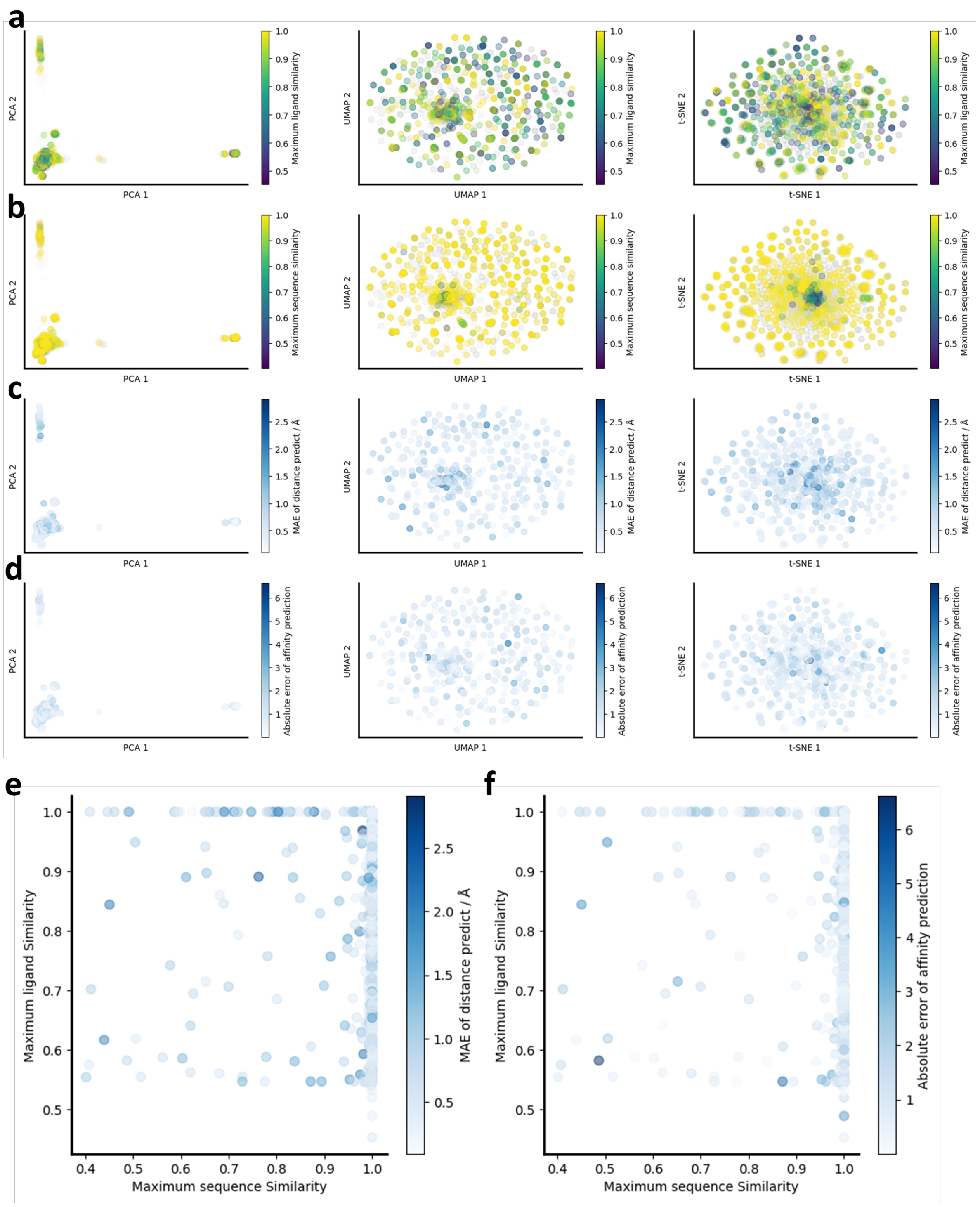


**Figure S4. Assessment of the capability of the model for generalisation. (a-d)** Visualizations of the complex feature space via PCA, t-SNE[7], and UMAP[7] (from left to right, respectively). The feature set for each complex was constructed by concatenating the Morgan fingerprints[8] (radius=2, dimension=1024) for ligands, with sequence features consist of: (i) 3-mer frequency, which counts the occurrences of each possible 3-length subsequence within the protein sequence, (ii) amino acid composition, and (iii) physico-chemical properties, including estimated hydrophobicity[9] and volume[10]. **(a)** Training set data points are shown in grey, with test set data points coloured according to ligand similarity to the training set, where dark purple indicates low similarity and yellow denotes high similarity. **(b)** Training data points are grey, with test set data points coloured by sequence similarity, transitioning from dark purple (low similarity) to yellow (high similarity). **(c)** Test set data points colored by the mean absolute error (MAE) of distance prediction, reflecting the average prediction error across all distance pairs in a complex. **(d)** Test set data points colored by the absolute error in p$K_d$ prediction. **(e)** Scatter plot correlating the MAE of distance prediction with both ligand and sequence similarity to the training set. **(f)** Scatter plot illustrating the relationship between affinity prediction error and similarity of ligand and sequence to the training set. Here, maximum ligand similarity represents the highest Tanimoto similarity score with any ligand in the training set, based on topological fingerprints in RDKit[11]. The maximum sequence similarity was calculated as the highest ratio of identical amino acids following global alignment without a gap penalty, as implemented in Biopython[12].

To demonstrate the ability of the model to predict distances when both the protein and the compound have not been seen in the training, we selected the DNA ligase A in complex with inhibitor 7-methoxy-6-methylpteridine-2,4-diamine as an example (PDB 4GLW). The sequence identity of this complex with the training set is below 0.41, while the Tanimoto similarity of the ligand with the training set is below 0.58. A comparison between the predicted and true distance matrices for PDB 4GLW revealed a MAE of 0.24 Å in prediction of the protein distance matrix ($\widehat{d}_{ij}$), 0.59 Å of ligand distance matrix ($\widehat{d}_{kl}$), and 0.83 Å of contact matrix ($\widehat{d}_{ik}$) (**Figure S5a,b**). The comparison between the estimated distance prediction error (**Figure S5c**) and the actual error (**Figure S5d**) indicates that the model is capable of assessing errors, although it tends to underestimate them. Furthermore, the predicted residues in contact with the ligand align closely with the actual binding pocket (**Figure S5e**), while the predicted atomic contacts with the pocket are in good agreement with the ground truth (**Figure S5f**). These results indicate that Ligand-Transformer is capable of accurately predicting contacts even when both the protein and the ligand have not been encountered in the training set.

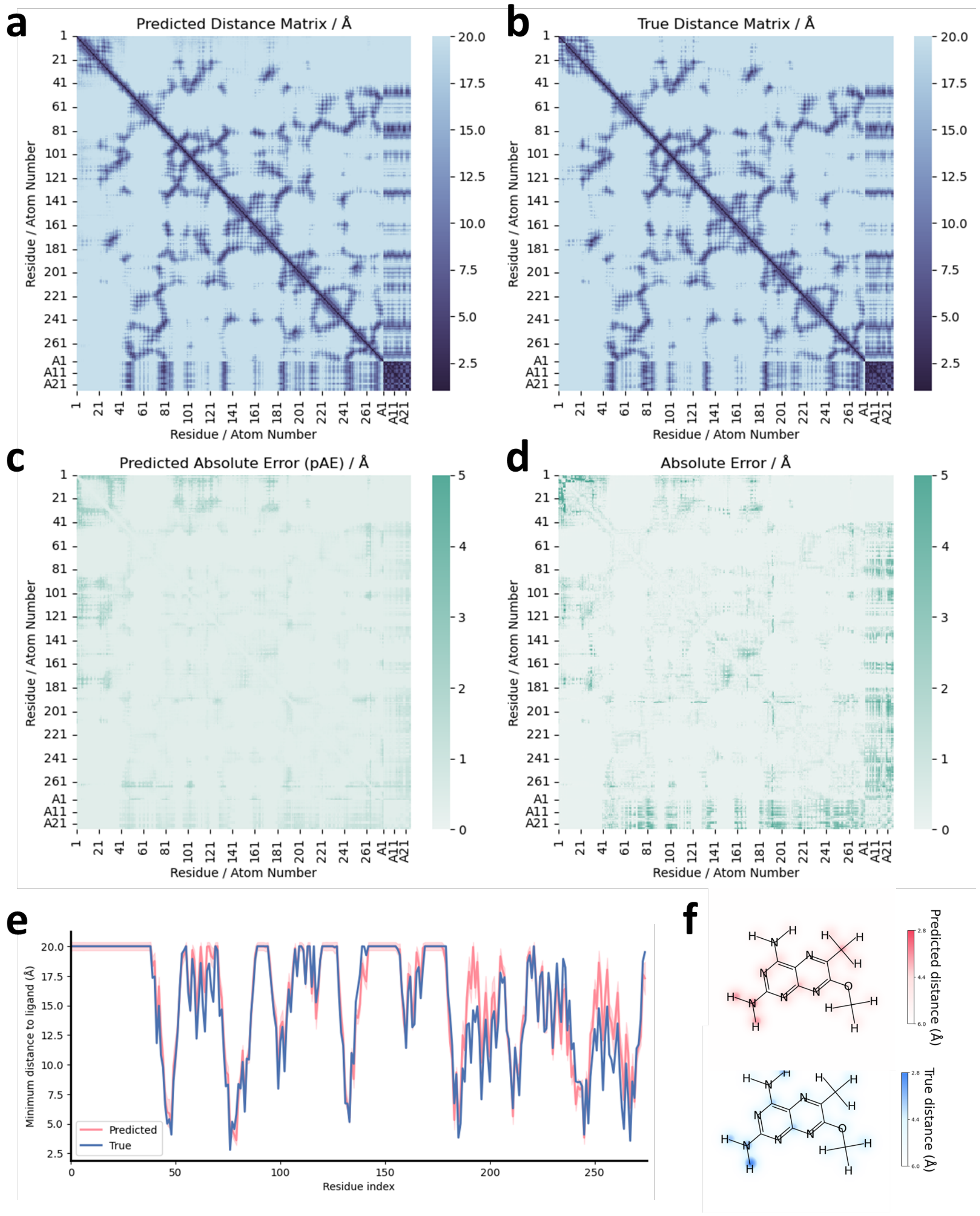


**Figure S5. Detailed distance prediction analysis for DNA ligase A complex with an inhibitor (PDB 4GLW).** **(a, b)** Heat maps displaying the predicted **(a)** and actual **(b)** distance matrices for DNA ligase A in complex with the inhibitor (ligand ID: 0XT). The axes represent residue numbers and atom numbers (prefixed with 'A') of the complex. **(c, d)** Heat maps showing the predicted **(c)** and actual **(d)** absolute errors in the distance matrices for PDB 4GLW. **(e)** Line graph comparing the nearest distances between the small molecule and the Cβ atom (or Cα for glycine) of each residue, providing a visualization of the binding region of the ligand within the protein. Predicted distances are marked with error bars derived from the predicted absolute error (pAE) as calculated by the confidence module (refer to **Algorithm 8**). **(f)** Contact heat map that indicates the proximity between each atom of the small molecule and the nearest Cβ atom (or Cα for glycine) of the protein.

### The Ligand-Transformer algorithm

We present the Ligand-Transformer algorithm in the form of a pseudocode. Scalar values are represented in italics, while vectors are represented in bold. Specifically, $\mathbf{n}$ represents a vector of nodes, and $\mathbf{e}$ represents a vector of edges. We use the indices $i$ and $j$ for the residues of a protein, and $k$ and $l$ for the atoms of a ligand, so that $(i, k)$ denotes an edge from residue $i$ of the protein to atom $k$ of the ligand. Although the edge representations are directional, for conciseness we use $\{\mathbf{e}_{ik}\}$ to represent the sum set of all $\{\mathbf{e}_{ik}\}$ and $\{\mathbf{e}_{ki}\}$, encompassing both edges from the residues of the protein to the atoms of the ligand and the edges from the atoms of the ligand to the residues of the protein. We use capitalized operator names when they encapsulate learned parameters. For functions without parameters, we use lowercase operator names. More details about the functions can be found in the following subsections. We use $\odot$ for element-wise multiplication, $\oplus$ for the outer sum, and $\mathbf{a}^{\top}\mathbf{b}$ for the dot product of two vectors.

#### Basic components and blocks

The following basic functions are utilized in the subsequent pseudocode:

**Linear:** Represents a linear transformation that involves multiplying the input by a weight matrix $W$ and adding a bias vector $\mathbf{b}$.

**LinearNoBias:** Denotes a linear transformation without the bias vector b. It involves only the multiplication of the input by the weight matrix $W$.

**LayerNorm:** This refers to layer normalization[13]. It operates on the channel dimensions and includes learnable per-channel gains and biases.

**ReLU:** This term denotes the Rectified Linear Unit (ReLU) activation function. It is a non-linear function that outputs the input directly if it is positive; otherwise, it outputs zero.

**Sigmoid:** This function denotes the sigmoid activation function that maps the input to a value in the range of 0 to 1, using the equation: $\text{sigmoid}(x) = 1/(1 + e^{-x})$.

$\mathbf{Softmax}_{ind}$**:** This function signifies the softmax activation function that normalizes a vector of numerical values into a probability distribution over the indices *ind*.

$\mathbf{Dropout}_{rate}$**:** This term designates the rate parameter of the dropout regularization technique. This method randomly nullifies a certain proportion ($rate$) of input units during training as a preventive measure against overfitting[14].

$\mathbf{Mean}_{ind}$**:** This function represents a process for calculating the mean value. It computes the average value across a specified index, $ind$.

$\mathbf{Concat}_{ind}$**:** This function is used for concatenating vectors based on the specified index, $ind$. If $ind$ is not provided, it defaults to concatenating two vectors into a single vector.

#### Transition block

The transition block facilitates non-linear transitions for the features of nodes or edges, while maintaining dimension invariance. A transition block comprises a layer normalization[13], which is applied to the input activations, succeeded by a standard 2-layer multilayer perceptron (MLP) utilizing ReLU as its activation function. The intermediate number of channels in the MLP expands the original number of channels by a factor of $n_{\text{expand}} = 4$. If we assume an input activation, denoted as a vector $\mathbf{a} \in \mathbb{R}^{c}$, where $c$ represents the dimension of $\mathbf{a}$, then the algorithm can be illustrated as:

**Algorithm 1** Transition block

**def** Transition $(\mathbf{a}, n_{\text{expand}} = 4)$:

$\mathbf{a}' = \text{Linear}(\mathbf{a})$ $\qquad$ $\mathbf{a}' \in \mathbb{R}^{c*n_{\text{expand}}}$

$\mathbf{a} \leftarrow \text{Linear}(\text{relu}(\mathbf{a}'))$

**return a**

---

#### Self-attention block

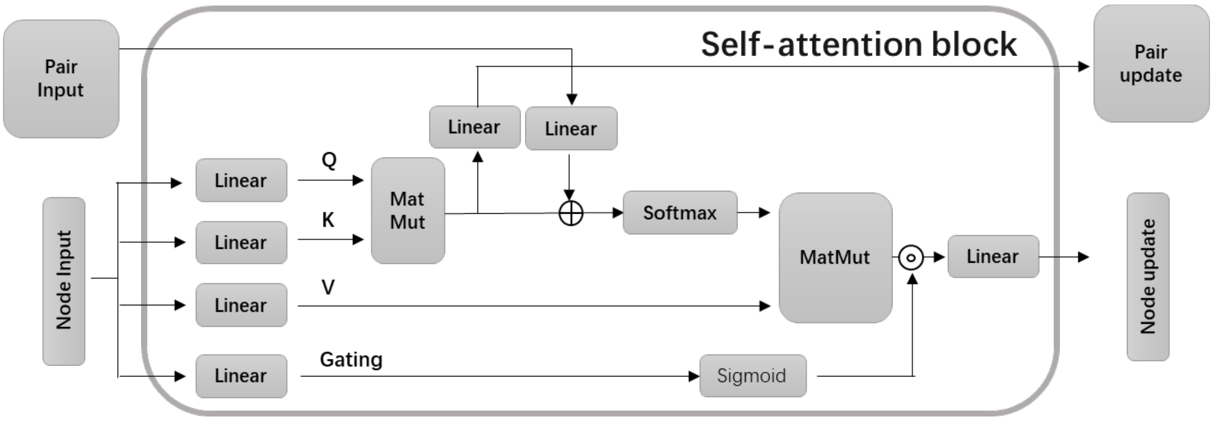


**Figure S6. Schematic representation of a gated self-attention block including pair bias and pair updating.**

The self-attention block is responsible for updating a graph representation consisting of both node features and edge features. It is a modified version of the basic self-attention[15] with pair bias[1], designed to facilitate effective communication between the edge features and node features. The attention block updates the edge features through the attention matrix derived from node features. The number of attention heads $N_{\text{head}}$ is a hyperparameter. Assuming $\{\mathbf{n}_i\}$ are features of dimension $c_{\text{n}}$ for each node, and $\{\mathbf{e}_{ij}\}$ donates features for each pair of nodes, $i$ and $j$, then the algorithm can be described as:

---

**Algorithm 2** Self-attention with pair bias

---

**def** SelfAttention ($\{\mathbf{n}_i\}, \{\mathbf{e}_{ij}\}, N_{\text{head}} = 8$):

*# Input projections*

$\mathbf{n}_i \leftarrow \text{LayerNorm}(\mathbf{n}_i)$

$\mathbf{e}_{ij} \leftarrow \text{LayerNorm}(\mathbf{e}_{ij})$

$c'_{\text{n}} = \dfrac{c_{\text{n}}}{N_{\text{head}}}$

$\mathbf{q}_i^h, \mathbf{k}_i^h, \mathbf{v}_i^h, = \text{LinearNoBias}(\mathbf{n}_i)$ $\qquad \mathbf{q}_i^{\text{h}}, \mathbf{k}_i^{\text{h}}, \mathbf{v}_i^{\text{h}} \in \mathbb{R}^c, h \in \{1, \dots, N_{\text{head}}\}$

$b_{ij}^h = \text{LinearNoBias}(\mathbf{e}_{ij})$

$\mathbf{g}_i^h = \text{sigmoid}(\text{Linear}(\mathbf{n}_i))$ $\qquad \mathbf{g}_i^{\text{h}} \in \mathbb{R}^{c'_{\text{n}}}$

*# Attention*

$a_{ij}^h = \dfrac{1}{\sqrt{c}} {\mathbf{q}_i^h}^{\top} \mathbf{k}_i^h$

*# Output edge features*

$\mathbf{e}_{ij} \leftarrow \text{Linear}\left(\text{relu}\left(\text{Linear}\left(\text{concat}_h\left(a_{ij}^h\right)\right)\right)\right)$

*# Output node features*

$a_{ij}^h \leftarrow \text{softmax}_{\text{j}}\left(a_{ij}^h + b_{ij}^h\right)$

$\mathbf{o}_i^h = \mathbf{g}_i^h \odot \sum_j a_{ij}^h \mathbf{v}_i^h$

$\mathbf{n}_i \leftarrow \text{Linear}\left(\text{concat}_h\left(\mathbf{o}_i^h\right)\right)$

**return** $\{\mathbf{n}_i\}, \{\mathbf{e}_{ij}\}$

---

### Protein encoder

The protein encoder takes the intermediate features from AlphaFold2, namely $\{\mathbf{f}_i^{\text{msa}}\}, \{\mathbf{f}_{ij}^{\text{pair}}\}$, and $\{\mathbf{f}_i^{\text{struc}}\}$, as input. These are transformed into initial representations for nodes $\{\mathbf{n}_i^{(0)}\}$ and representations for edges as pairs of residues $\{\mathbf{e}_{ij}^{(0)}\}$ in the Ligand-Transformer model. Within this model, the node dimension ($c_\text{n}$) is set to 384, while the edge dimension ($c_\text{e}$) is set to 128.

**Algorithm 3** Encoding input protein representations from AlphaFold2

**def** ProteinEncoder $(\{\mathbf{f}_i^{\text{msa}}\}, \{\mathbf{f}_{ij}^{\text{pair}}\}, \{\mathbf{f}_i^{\text{struc}}\})$:

*# Input projections*

$\mathbf{f}_i^{\text{msa}} \leftarrow \text{LayerNorm}(\text{Transition}(\mathbf{f}_i^{\text{msa}}))$

$\mathbf{f}_{ij}^{\text{pair}} \leftarrow \text{LayerNorm}(\text{Transition}(\mathbf{f}_{ij}^{\text{pair}}))$

$\mathbf{f}_i^{\text{struc}} \leftarrow \text{LayerNorm}(\text{Transition}(\mathbf{f}_i^{\text{struc}}))$

*# Concatenate residue-wise features*

$\mathbf{f}_i^{\text{resi}} = \text{concat}(\mathbf{f}_i^{\text{msa}}, \mathbf{f}_i^{\text{struc}})$ $\quad \mathbf{f}_i^{\text{resi}} \in \mathbb{R}^{c_{\text{msa}}+c_{\text{struct}}}$

*# Output projection*

$\mathbf{e}_{ij}^{(0)} \leftarrow \text{Transition}(\mathbf{f}_{ij}^{\text{pair}})$ $\quad \mathbf{e}_{ij}^{(0)} \in \mathbb{R}^{c_\text{e}}$

$\mathbf{n}_i^{(0)} \leftarrow \text{Transition}(\mathbf{f}_i^{\text{resi}})$ $\quad \mathbf{n}_i^{(0)} \in \mathbb{R}^{c_\text{n}}$

**return** $\{\mathbf{n}_i^{(0)}\}, \{\mathbf{e}_{ij}^{(0)}\}$

### Ligand encoder

The ligand encoder transforms the molecular representations output by GraphMVP[2] into representations within the Ligand-Transformer model. The GraphMVP representation defines the molecular graph as a directed sparse graph, meaning that bond feature $\mathbf{f}_{kl}^{\text{bond}}$ only exists between bonded atoms $(k, l)$. However, in Ligand-Transformer, we represent the molecule as a complete graph, meaning we maintain edge features $\mathbf{e}_{kl}$ between each pair of atoms $(k, l)$. Thus, during the encoding process, we perform padding to complete the edge features. The information is then updated through four layers of self-attention blocks.

**Algorithm 4** Encoding input ligand representations from GraphMVP

**def** LigandEncoder $(\{\mathbf{f}_k^{\text{atom}}\}, \{\mathbf{f}_{kl}^{\text{bond}}\})$:

*# Padding to complete graph*

$\mathbf{f}_{kl}^{\text{atom\_pair}} = \mathbf{f}_{kl}^{\text{bond}}$ if bond $(k, l)$ exists else $\mathbf{0}$

*# Input projections*

$\mathbf{f}_k^{\text{atom}} \leftarrow \text{Transition}\ (\mathbf{f}_k^{\text{atom}})$

$\mathbf{f}_{kl}^{\text{atom\_pair}} \leftarrow \text{Transition}\ (\mathbf{f}_{kl}^{\text{atom\_pair}})$

*# Communication between atom representations and edge representations*

**for all** $m \in [1, \ldots, 4]$ *# not sharing weights*

$\quad \{\mathbf{f}_k^{\text{atom}}\}, \{\mathbf{f}'^{\text{bond}}_{kl}\} \mathrel{+}= \text{dropout}_{drop\_rate_{\text{attention}}}(\text{SelfAttention}\ (\{\mathbf{f}_k^{\text{atom}}\}, \{\mathbf{f}_{kl}^{\text{bond}}\}, N_{\text{head}} = 6)$

$\quad \mathbf{f}_k^{\text{atom}} \mathrel{+}= \text{dropout}_{drop\_rate_{\text{transition}}}(\text{Transition}(\text{LayerNorm}(\mathbf{f}_k^{\text{atom}})))$

$\quad \mathbf{f}_{kl}^{\text{bond}} \mathrel{+}= \text{dropout}_{drop\_rate_{\text{transition}}}\left(\text{Transition}\left(\text{LayerNorm}(\mathbf{f}_{kl}^{\text{bond}})\right)\right)$

*# Output projections*

$\mathbf{e}_{kl}^{(0)} = \text{Linear}(\text{LayerNorm}(\mathbf{f}_{kl}^{\text{bond}}))$ $\quad \mathbf{e}_{kl}^{(0)} \in \mathbb{R}^{c_\text{e}}$

$\mathbf{n}_k^{(0)} = \text{Linear}(\text{LayerNorm}(\mathbf{f}_k^{\text{atom}}))$ $\quad \mathbf{n}_k^{(0)} \in \mathbb{R}^{c_\text{n}}$

**return** $\{\mathbf{n}_k^{(0)}\}, \{\mathbf{e}_{kl}^{(0)}\}$

### Cross-modal attention module

After the separate encoding of the protein and ligand, all node representations, whether they are residues or atoms, have the same dimension $c_{\mathrm{n}}$. Similarly, all edge representations, whether they are pairs of residues or pairs of atoms, also share the same vector dimension $c_{\mathrm{e}}$. Despite the protein-ligand being a heterogeneous graph, we do not explicitly differentiate between the types of edges and nodes in the cross-modal attention module. The symbol $\{\mathbf{n}^{(m)}\}$ represents the set of all residues and atom nodes, while $\{\mathbf{e}^{(m)}\}$ represents the collection of all pairs of residues, atom pairs, and residue-atom pairs. Interaction between edge information and node information is carried out via a self-attention block, each followed by a transition layer for nonlinear transformation. The output of each layer is added to the current representations via a residual connection[16] and passed through a Dropout[14] layer before they are added together.

**Algorithm 5** Update node representations and edge representations

**def** CrossModalAttention $\left(\{\mathbf{n}_i^{(m)}\}, \{\mathbf{n}_k^{(m)}\}, \{\mathbf{e}_{ij}^{(m)}\}, \{\mathbf{e}_{kl}^{(m)}\}, \{\mathbf{e}_{ik}^{(m)}\}\right)$:

*# The type of nodes and edges are not explicitly distinguished in this block*

$\{\mathbf{n}^{(m)}\} \triangleq \{\mathbf{n}_i^{(m)}\} + \{\mathbf{n}_k^{(m)}\}$

$\{\mathbf{e}^{(m)}\} \triangleq \{\mathbf{e}_{ij}^{(m)}\} + \{\mathbf{e}_{kl}^{(m)}\} + \{\mathbf{e}_{ik}^{(m)}\}$

*# Communication and transition*

$\{\mathbf{n}^{(m)}\}, \{\mathbf{e}^{(m)}\} \mathrel{+}= \mathrm{dropout}_{drop\_rate_{\mathrm{attention}}}(\mathrm{SelfAttention}\ (\{\mathbf{n}^{(m)}\}, \{\mathbf{e}^{(m)}\}))$

$\mathbf{n}^{(m+1)} = \mathrm{dropout}_{drop\_rate_{\mathrm{transition}}}\left(\mathrm{Transition}\left(\mathrm{LayerNorm}\left(\mathbf{n}^{(m)}\right)\right)\right) + \mathbf{n}^{(\mathrm{m})}$

$\mathbf{e}^{(m+1)} = \mathrm{dropout}_{drop\_rate_{\mathrm{transition}}}\left(\mathrm{Transition}\left(\mathrm{LayerNorm}\left(\mathbf{e}^{(m)}\right)\right)\right) + \mathbf{e}^{(m)}$

**return** $\{\mathbf{n}_i^{(m+1)}\}, \{\mathbf{n}_k^{(m+1)}\}, \{\mathbf{e}_{ij}^{(m+1)}\}, \{\mathbf{e}_{kl}^{(m+1)}\}, \{\mathbf{e}_{ik}^{(m+1)}\}$

### Affinity prediction head

After passing through twelve layers of cross-modal attention modules, we obtain the final representations of the protein residues $\{\mathbf{n}_i\}$ and the small molecule atoms $\{\mathbf{n}_k\}$. We apply average pooling separately to these two sets of representations, and then concatenate them to obtain a joint representation vector of the complex system $\hat{\mathbf{n}}$, which is used for predicting binding affinity. During the training phase, we employ the mean squared error (MSE) to calculate the discrepancy in affinity prediction.

**Algorithm 6** Predict binding affinity

**def** AffinityHead $(\{\mathbf{n}_i\}, \{\mathbf{n}_k\}, true\_affinity)$:

*# Input projection*

$\mathbf{n}_i \leftarrow \mathrm{Linear}(\mathrm{relu}(\mathrm{Linear}(\mathrm{dropout}_{drop\_rate_{\mathrm{affinity}}}(\mathrm{LayerNorm}(\mathbf{n}_i)))))$

$\mathbf{n}_k \leftarrow \mathrm{Linear}(\mathrm{relu}(\mathrm{Linear}(\mathrm{dropout}_{drop\_rate_{\mathrm{affinity}}}(\mathrm{LayerNorm}(\mathbf{n}_k)))))$

*# Average pooling*

$\hat{\mathbf{n}}^{\mathrm{protein}} = \mathrm{mean}_i(\{\mathbf{n}_i\})$

$\hat{\mathbf{n}}^{\mathrm{ligand}} = \mathrm{mean}_k(\{\mathbf{n}_k\})$

$\hat{\mathbf{n}} = \mathrm{concat}\left(\hat{\mathbf{n}}^{\mathrm{protein}}, \hat{\mathbf{n}}^{\mathrm{ligand}}\right)$

*# Output projection*

$affinity = \mathrm{Linear}(\mathrm{relu}(\mathrm{Linear}(\mathrm{relu}(\mathrm{Linear}(\hat{\mathbf{n}})))))$

*# Calculate loss when training*

$\mathcal{L}_{\mathrm{affinity}} = (affinity - true\_affinity)^2$

**return** $affinity, \mathcal{L}_{\mathrm{affinity}}$

### Distance prediction head

The distance prediction head initially transforms the pair representations and categorizes them into a set of distance bins. These bins, denoted as $\mathbf{v}_{\text{bins}}$, vary across different models as detailed below. The specific bin definition used in each model can be found in **Tables S2-S4**. The model then calculates the probabilities using a softmax function. For prediction targets, we utilize one-hot encoded binned values from the ground-truth distances, and the loss is computed as the cross-entropy loss averaged over all these distances. The model determines three types of distances. The first is the distance between residues $i$ and $j$, $\widehat{d}_{ij}$, which is defined as the distance between their Cβ atoms, unless the residue is glycine, in which case the Cα atom distance is used. The second type of distance, $\widehat{d}_{kl}$, measures the distance between atoms $k$ and $l$ of the ligand. Lastly, the distance between residue $i$ of the protein and an atom $k$ of the ligand, $\widehat{d}_{ik}$, is calculated based on the distance from the Cβ atoms (or the Cα atom for glycine) to the atom of the ligand. Each of these distances is calculated independently using their respective edge representations, with the parameters not sharing across different edge types.

**Algorithm 7** Predict distance

**def** DistanceHead $\left(\{\mathbf{e}_{ij}\}, \{\mathbf{e}_{kl}\}, \{\mathbf{e}_{ik}\}, \{d_{ij}^{\text{true}}\}, \{d_{kl}^{\text{true}}\}, \{d_{ik}^{\text{true}}\}\right)$:

*# Linear projection*

$\mathbf{p}_{ij} = \text{Linear}(\mathbf{e}_{ij})$

$\mathbf{p}_{kl} = \text{Linear}(\mathbf{e}_{kl})$

$\mathbf{p}_{ik} = \text{Linear}(\mathbf{e}_{ik})$ $\qquad \mathbf{p}_{ij}, \mathbf{p}_{kl}, \mathbf{p}_{ik} \in \mathbb{R}^{|\mathbf{v}_{\text{bins}}|}$

*# Symmetrize the logit values*

$\mathbf{p}_{ij} \leftarrow \text{softmax}(\mathbf{p}_{ij} + \mathbf{p}_{ji})$

$\mathbf{p}_{kl} \leftarrow \text{softmax}(\mathbf{p}_{kl} + \mathbf{p}_{lk})$

$\mathbf{p}_{ik} \leftarrow \text{softmax}(\mathbf{p}_{ik} + \mathbf{p}_{ki})$

*# Predicted distance is the expected value of the distribution*

$\widehat{d}_{ij} = \mathbf{p}_{ij}^{\top}\mathbf{v}_{\text{bins}}$

$\widehat{d}_{kl} = \mathbf{p}_{kl}^{\top}\mathbf{v}_{\text{bins}}$

$\widehat{d}_{ik} = \mathbf{p}_{ik}^{\top}\mathbf{v}_{\text{bins}}$

*# Calculate loss when training*

$$\begin{aligned}\mathcal{L}_{\text{distance}} = \ & \text{mean}_{(i,j)}\left(\text{one\_hot}\left(d_{ij}^{\text{true}}, \mathbf{v}_{\text{bins}}\right)^{\top} \log \mathbf{p}_{ij}\right) \\ & + \text{mean}_{(k,l)}\left(\text{one\_hot}(d_{kl}^{\text{true}}, \mathbf{v}_{\text{bins}})^{\top} \log \mathbf{p}_{kl}\right) \\ & + \text{mean}_{(i,k)}\left(\text{one\_hot}(d_{ik}^{\text{true}}, \mathbf{v}_{\text{bins}})^{\top} \log \mathbf{p}_{ik}\right)\end{aligned}$$

*# Calculate absolute error when training*

$err_{ij} = |\widehat{d}_{ij} - d_{ij}^{\text{true}}|$

$err_{kl} = |\widehat{d}_{kl} - d_{kl}^{\text{true}}|$

$err_{ik} = |\widehat{d}_{ik} - d_{ik}^{\text{true}}|$

**return** $\widehat{d}_{ij}, \widehat{d}_{kl}, \widehat{d}_{ik}, \mathcal{L}_{\text{distance}}, err_{ij}, err_{kl}, err_{ik}$

### Distance error prediction

We estimate the confidence of our distance predictions by predicting the absolute error. Following a similar approach to the distance prediction head, we transform the pair representation into 100 bins via a linear transformation. These bins range uniformly from 0.5 to 15.5 Å, and the model assigns a probability for the prediction error to fall within each bin. During model training, we first obtain the prediction error from the distance prediction head. This error is then converted into a label via one-hot encoding, and the loss is computed using cross-entropy.

**Algorithm 8** Distance error prediction

**def** DistanceErrorHead $\left(\{\mathbf{e}_{ij}\}, \{\mathbf{e}_{kl}\}, \{\mathbf{e}_{ik}\}, err_{ij}, err_{kl}, err_{ik}\right)$:

*# Linear projection*

$\mathbf{p}_{ij} = \mathrm{Linear}(\mathbf{e}_{ij})$

$\mathbf{p}_{kl} = \mathrm{Linear}(\mathbf{e}_{kl})$

$\mathbf{p}_{ik} = \mathrm{Linear}(\mathbf{e}_{ik})$ $\mathbf{p}_{ij}, \mathbf{p}_{kl}, \mathbf{p}_{ik} \in \mathbb{R}^{|\mathbf{v}_{\mathrm{error\_bins}}|}$

*# Symmetrize the logit values*

$\mathbf{p}_{ij} \leftarrow \mathrm{softmax}(\mathbf{p}_{ij} + \mathbf{p}_{ji})$

$\mathbf{p}_{kl} \leftarrow \mathrm{softmax}(\mathbf{p}_{kl} + \mathbf{p}_{lk})$

$\mathbf{p}_{ik} \leftarrow \mathrm{softmax}(\mathbf{p}_{ik} + \mathbf{p}_{ki})$

*# Predicted distance error is the expected value of the distribution*

$\widehat{err}_{ij} = {\mathbf{p}_{ij}}^{\top} \mathbf{v}_{\mathrm{error\_bins}}$

$\widehat{err}_{kl} = {\mathbf{p}_{kl}}^{\top} \mathbf{v}_{\mathrm{error\_bins}}$

$\widehat{err}_{ik} = {\mathbf{p}_{ik}}^{\top} \mathbf{v}_{\mathrm{error\_bins}}$

*# Calculate loss when training*

$$\begin{aligned}\mathcal{L}_{\mathrm{distance\_error}} = \ & \mathrm{mean}_{(i,j)}\left(\mathrm{one\_hot}\left(err_{ij}, \mathbf{v}_{\mathrm{error\_bins}}\right)^{\top} \log \mathbf{p}_{ij}\right) \\ & + \mathrm{mean}_{(k,l)}\left(\mathrm{one\_hot}\left(err_{kl}, \mathbf{v}_{\mathrm{error\_bins}}\right)^{\top} \log \mathbf{p}_{kl}\right) \\ & + \mathrm{mean}_{(i,k)}\left(\mathrm{one\_hot}\left(err_{ik}, \mathbf{v}_{\mathrm{error\_bins}}\right)^{\top} \log \mathbf{p}_{ik}\right)\end{aligned}$$

**return** $\widehat{err}_{ij}, \widehat{err}_{kl}, \widehat{err}_{ik}, \mathcal{L}_{\mathrm{distance\_error}}$

### Ligand-Transformer

Here we introduce the complete architecture of Ligand-Transformer. We initially encode the inputs from both the protein and small molecules separately, leading to a unified complete graph representation of them, namely $\left\{\mathbf{n}_i^{(0)}\right\}$, $\left\{\mathbf{n}_k^{(0)}\right\}$, $\left\{\mathbf{e}_{ij}^{(0)}\right\}$, and $\left\{\mathbf{e}_{kl}^{(0)}\right\}$. For all protein-small molecule contact edges, represented as $\left\{\mathbf{e}_{ik}^{(0)}\right\}$, we set them as zero vectors. These vectors are padded to form a protein-small molecule complete graph. Following this, we utilize twelve layers of non-parameter-shared cross-modal attention modules to facilitate information exchange and updates between nodes and edges. The resulting outputs are highly processed representations of the complex system. The node representations are used as input for the affinity prediction head for activity prediction, while edge representations feed into the distance prediction modules.

**Algorithm 9** Ligand-Transformer main architecture

**def** Ligand-Transformer $\left(\{\mathbf{f}_i^{\mathrm{msa}}\}, \{\mathbf{f}_{ij}^{\mathrm{pair}}\}, \{\mathbf{f}_i^{\mathrm{struc}}\}, \{\mathbf{f}_k^{\mathrm{atom}}\}, \{\mathbf{f}_{kl}^{\mathrm{bond}}\}\right)$:

*# Input encoding*

$\left\{\mathbf{n}_i^{(0)}\right\}, \left\{\mathbf{e}_{ij}^{(0)}\right\} = \mathrm{ProteinEncoder}\left(\{\mathbf{f}_i^{\mathrm{msa}}\}, \{\mathbf{f}_{ij}^{\mathrm{pair}}\}, \{\mathbf{f}_i^{\mathrm{struc}}\}\right)$

$\left\{\mathbf{n}_k^{(0)}\right\}, \left\{\mathbf{e}_{kl}^{(0)}\right\} = \mathrm{LigandEncoder}\left(\{\mathbf{f}_k^{\mathrm{atom}}\}, \{\mathbf{f}_{kl}^{\mathrm{bond}}\}\right)$

$\left\{\mathbf{e}_{ik}^{(0)}\right\} = \mathbf{0}$ $\mathbf{e}_{ik}^{(0)} \in \mathbb{R}^{c_{\mathrm{e}}}$

*# Update representations*

**for all** $m \in [1, \ldots, 12]$ *# not sharing weights*

$$\begin{aligned}&\left\{\mathbf{n}_i^{(m)}\right\}, \left\{\mathbf{n}_k^{(m)}\right\}, \left\{\mathbf{e}_{ij}^{(m)}\right\}, \left\{\mathbf{e}_{kl}^{(m)}\right\}, \left\{\mathbf{e}_{ik}^{(m)}\right\} \leftarrow \\ &\quad \mathrm{CrossModalAttention}\left(\left\{\mathbf{n}_i^{(m-1)}\right\}, \left\{\mathbf{n}_k^{(m-1)}\right\}, \left\{\mathbf{e}_{ij}^{(m-1)}\right\}, \left\{\mathbf{e}_{kl}^{(m-1)}\right\}, \left\{\mathbf{e}_{ik}^{(m-1)}\right\}\right)\end{aligned}$$

*# Predict affinity and distance*

$affinity, \mathcal{L}_{\mathrm{affinity}} = \mathrm{AffinityHead}\left(\left\{\mathbf{n}_i^{(12)}\right\}, \left\{\mathbf{n}_k^{(12)}\right\}, true\_affinity\right)$

$$\begin{aligned}&\hat{d}_{ij}, \hat{d}_{kl}, \hat{d}_{ik}, \mathcal{L}_{\mathrm{distance}}, err_{ij}, err_{kl}, err_{ik} \\ &\quad = \mathrm{DistanceHead}\left(\left\{\mathbf{e}_{ij}^{(12)}\right\}, \left\{\mathbf{e}_{kl}^{(12)}\right\}, \left\{\mathbf{e}_{ik}^{(12)}\right\}, \{d_{ij}^{\mathrm{true}}\}, \{d_{kl}^{\mathrm{true}}\}, \{d_{ik}^{\mathrm{true}}\}\right)\end{aligned}$$

*# Predict distance prediction errors*

$$\widehat{err}_{ij}, \widehat{err}_{kl}, \widehat{err}_{ik}, \mathcal{L}_{\text{distance\_error}} = \text{DistanceErrorHead}\left(\{\mathbf{e}_{ij}\}, \{\mathbf{e}_{kl}\}, \{\mathbf{e}_{ik}\}, err_{ij}, err_{kl}, err_{ik}\right)$$

*# Summing losses when training*

$$\mathcal{L} = \mathcal{L}_{\text{affinity}} * w_{\text{affinity}} + \mathcal{L}_{\text{distance}} * w_{\text{distance}} + \mathcal{L}_{\text{distance\_error}} * w_{\text{distance\_error}}$$

**return** $affinity, \hat{d}_{ij}, \hat{d}_{kl}, \hat{d}_{ik}, \mathcal{L}, \widehat{err}_{ij}, \widehat{err}_{kl}, \widehat{err}_{ik}$

### Training

#### *Training details*

The models used in different tasks have the same framework, but slightly different training hyperparameters. Model A, which has undergone hyperparameter optimization and exhibits the best performance in activity prediction and distance prediction, is used for comparison with other baseline models and also for calculating the conformational selectivity of ABL inhibitors. The screening of $\text{EGFR}^{\text{LTC}}$ inhibitors is done using Model B, and the fine-tuning on the $\text{EGFR}^{\text{LTC}}$-290 dataset is also based on this model. The screening of Aβ42 inhibitors is conducted using Model C. All models are trained using the Adam optimizer[17]. All training runs were performed on 4 NVIDIA A100-SXM-80GB GPUs with 2 AMD EPYC 7763 64-Core Processor 1.8 GHz and 1000 GiB RAM.

#### *Training of the hyperparameters*

***Model A.*** The training of Model A is carried out in three steps. First, we conducted warm-up training with 4,480 complexes (PDBbind_4k) that only included single chains with lengths less than 384. During this training, we incorporated a warm-up strategy where the learning rate linearly increases, followed by a linear decrease in both the learning rate and $w_{\text{distance}}$. This training yielded Model A.0. Next, we proceeded to train with the complete training set (PDBbind_10k) of 10,375 data points. This step produced Model A.1, where both the learning rate and $w_{\text{distance}}$ also decreased linearly throughout the training process. Finally, after adding the Distance error prediction module, we fine-tuned the model to produce the final version of Model A. In Model A, $\mathbf{v}_{\text{bins}}$ in the distance prediction head employs a set of 100 distance bins that stretch from 1 to 20 Å, with each bin sharing the same width. In **Table S1** we summarize the training hyperparameters of Model A.

**Table S2.** Training details of Model A.

| Model ID | A.0 | A.1 | A |
|---|---|---|---|
| Parameters initialized from | Random | A.0 | A.1 |
| Training dataset | PDBbind_4k | PDBbind_10k | PDBbind_10k |
| Learning rate | 5.4e-4 to 3e-4 | 3e-4 to 1e-4 | 1e-4 |
| Warm-up epochs | 5 | 0 | 0 |
| $w_{\text{affinity}}$ | 0.5 | 0.5 | 0.5 |
| $w_{\text{distance}}$ | 50 to 2.5 | 2.5 to 1.0 | 1.0 |
| $w_{\text{distance\_error}}$ | 0.0 | 0.0 | 0.5 |
| $drop\_rate_{\text{attention}}$ | 0.45 | 0.45 | 0.45 |
| $drop\_rate_{\text{transition}}$ | 0.15 | 0.15 | 0.15 |
| $drop\_rate_{\text{affinity}}$ | 0.06 | 0.06 | 0.06 |
| Batch_size | 8 | 8 | 8 |
| Number of epochs | 260 | 60 | 10 |

***Model B.*** In training Model B, we first trained on PDBbind_10k to obtain Model B.0. Since Model B was not employed for evaluating performance on PDBbind, we maximized the utilization of available datasets by training all data points in PDBbind2020-subset, adding the distance error prediction head on the basis of Model B.0 to derive Model B. We performed a 10-fold cross-validation on the $\text{EGFR}^{\text{LTC}}$-290 dataset, equally dividing it into 10 parts, and fine-tuned Model B to obtain Model FT1 to FT10. In Model B and Model FT1 to FT10, $\mathbf{v}_{\text{bins}}$ in the distance prediction head uses a set of 64 distance bins stretching from 2.3125 Å to 21.6875 Å, with each bin sharing the same width. This distance definition is identical to that used in AlphaFold2[1]. In **Table S2** we summarize the training hyperparameters of Model B and fine-tuned models on $\text{EGFR}^{\text{LTC}}$-290.

**Table S3.** Model B and fine-tuned models on $EGFR^{LTC}$-290.

| Model ID | B.0 | B | FT1 to FT10 |
|---|---|---|---|
| Parameters initialized from | Random | B.0 | B |
| Training dataset | PDBbind_10k | PDBbind2020-subset | $EGFR^{LTC}$-290 (ten-fold cross-validation) |
| Learning rate | 5e-5 | 4e-5 | 2e-7 |
| Warm-up epochs | 0 | 0 | 0 |
| $w_{\text{affinity}}$ | 0.5 | 0.5 | 0.5 |
| $w_{\text{distance}}$ | 50 | 50 | 0.0 |
| $w_{\text{distance\_error}}$ | 0.0 | 0.5 | 0.0 |
| $drop\_rate_{\text{attention}}$ | 0.3 | 0.3 | 0.65 |
| $drop\_rate_{\text{transition}}$ | 0.0 | 0.0 | 0.5 |
| $drop\_rate_{\text{affinity}}$ | 0.0 | 0.0 | 0.0 |
| Batch_size | 8 | 8 | 4 |
| Number of epochs | 36 | 16 | 100 |

***Model C.*** In training Model C, we only used the PDBbind_4k dataset. In Model C, $\mathbf{v}_{\text{bins}}$ are not evenly divided. There were 100 bins in total: the first 50 bins stretch from 1 Å to 7 Å, with a uniform interval, and the remaining 50 bins extend from 7 to 20 Å, also with a uniform interval. In **Table S3** we summarize the training hyperparameters of Model C.

**Table S4.** Model C and fine-tuned models on $EGFR^{LTC}$-290.

| Model ID | C |
|---|---|
| Parameters initialized from | Random |
| Training dataset | PDBbind_4k |
| Learning rate | 1e-5 |
| Warm-up epochs | 0 |
| $w_{\text{affinity}}$ | 0.5 |
| $w_{\text{distance}}$ | 0.5 |
| $w_{\text{distance\_error}}$ | 0.0 |
| $drop\_rate_{\text{attention}}$ | 0.15 |
| $drop\_rate_{\text{transition}}$ | 0.1 |
| $drop\_rate_{\text{affinity}}$ | 0.0 |
| Batch_size | 8 |
| Number of epochs | 250 |

#### *Ablation test*

***Training details of ablation test.*** Without AlphaFold2 embedding: For model without the protein representations generated by AlphaFold2, we replace $\{\mathbf{f}_i^{\text{msa}}\}$ with $\{\mathbf{f}_i^{\text{aa\_type}}\}$, where $\mathbf{f}_i^{\text{aa\_type}}$ is one-hot coded vector of the amino acid type of residue $i$; we replace $\{\mathbf{f}_{ij}^{\text{pair}}\}$ with $\{\mathbf{f}_{ij}^{\text{pair\_pos}}\}$, where $\mathbf{f}_{ij}^{\text{pair\_pos}}$ is the one-hot binned position number of residue $i$ minus the position number of residue $j$ in the protein sequence, with the minimum cut-off of -63 and maximum cut-off of 63. $\{\mathbf{f}_i^{\text{struc}}\}$ are not used in the model trained without AlphaFold2 embedding.

Without GraphMVP embedding: For model without the protein representations generated by AlphaFold2, we replace $\{\mathbf{f}_k^{\text{atom}}\}$ with $\{\mathbf{f}_k^{\text{atom\_type}}\}$, where $\mathbf{f}_k^{\text{atom\_type}}$ is one-hot coded vector of the element of atom $k$; we replace $\{\mathbf{f}_{kl}^{\text{bond}}\}$ with $\{\mathbf{f}_{kl}^{\text{bond\_type}}\}$, where $\mathbf{f}_{kl}^{\text{bond\_type}}$ is the one-hot coded vector representing the bond type between atom $k$ and $l$. The training hyperparameters for ablation test are the same as Model A.

#### *Ablation test result*

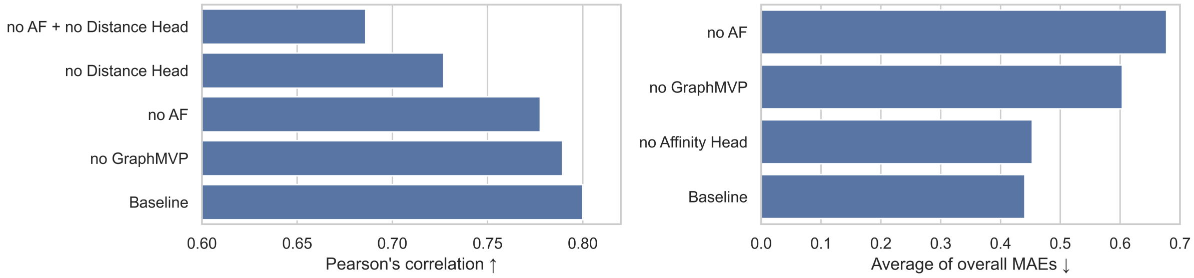


**Figure S7. Ablation study showing the impact of pretrained protein and ligand representations and joint training on model performance.** The figure illustrates the decrease in model performance upon the removal of pretrained embeddings and joint training for both affinity and distance prediction tasks. The affinity prediction is evaluated using Pearson's correlation between the predicted and measured p$K_d$ values. For distance prediction, the evaluation metric is the average of the overall MAEs of distance matrix.

To evaluate the impact of using pretrained representations of protein and ligand, as well as the joint training of affinity and distance prediction tasks, we carried out ablation experiments comparing the performance of the model without these components. The omission of pretrained protein and ligand representations resulted in performance decreases for both affinity and distance prediction tasks (**Figure S7**). Specifically, the Pearson's correlation for affinity prediction decreased to 0.78 and 0.77, and the average of overall MAEs of distance matrix – where it represents the MAE of the entire distance matrix for each complex – increased to 0.67 and 0.60, when we omitted the embeddings from AlphaFold2 and GraphMVP, respectively. These results illustrate that the pretrained representations of proteins and small molecules were beneficial for activity prediction and, particularly, distance prediction.

Furthermore, when joint training of affinity and distance prediction was removed, the Pearson's correlation for affinity prediction decreased to 0.74. In particular, the removal of both the AlphaFold2 embedding and the distance prediction resulted in a substantial decrease in the Pearson's correlation for affinity prediction, dropping it to 0.68. This finding suggests that the structure information integrated during training is important for activity prediction. In contrast, when we predicted distance alone, the accuracy did not decline significantly, implying that the activity information did not provide substantial additional information for distance prediction.

### Dataset preparation

***Protein sequence truncation.*** If the length of the sequence is less than or equal to 384, we retained the entire sequence. When truncation is necessary, we first calculated the minimum distance of each protein chain to the ligand, which is defined as the smallest distance among all the atoms in each chain and all the atoms in the ligand. We discarded all amino acids in the chains that are more than 8 Å away from the ligand. If the sequence length still exceeds 384, then we used Pfam[18] to identify the domains of the protein based on the sequence. Subsequently, we computed the minimal distance of all the domains to the ligand. This distance was defined as the smallest distance between all the atoms in the domain and all the atoms in the ligand. We progressively deleted domains that are more than 8 Å distant from the ligand until the sequence length is reduced to 384 or less.

***Dataset split.*** We randomly partitioned the PDBbind2020-subset into training, validation, and testing sets, which included 10,375, 640, and 936 data points respectively. The distributions of the training, validation, and test sets are alike, indicating that there is no bias in the dataset division (**Figure S8**).

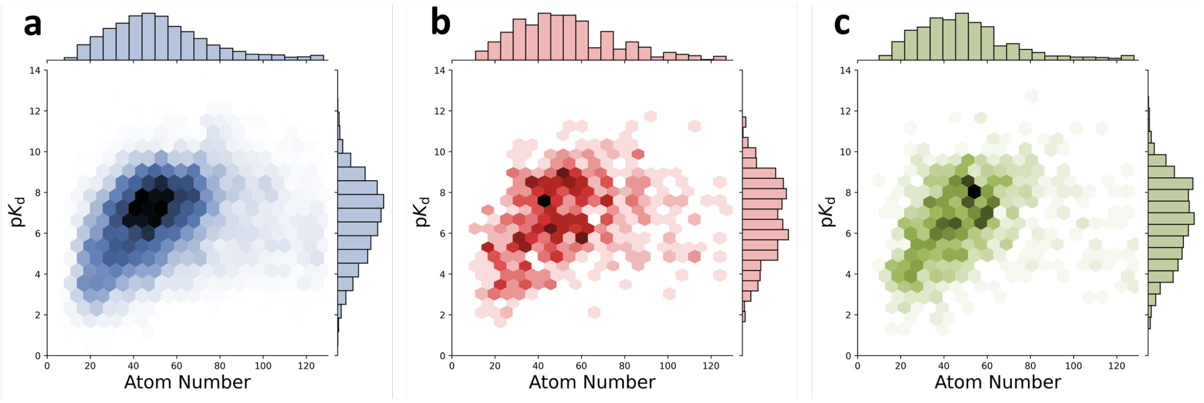


**Figure S8. Distribution of atom numbers of ligands and binding constants.** (**a**) Training set, (**b**) Validation set, and (**c**) Test set.

#### Fine-tuning on the EGFR$^{LTC}$ dataset

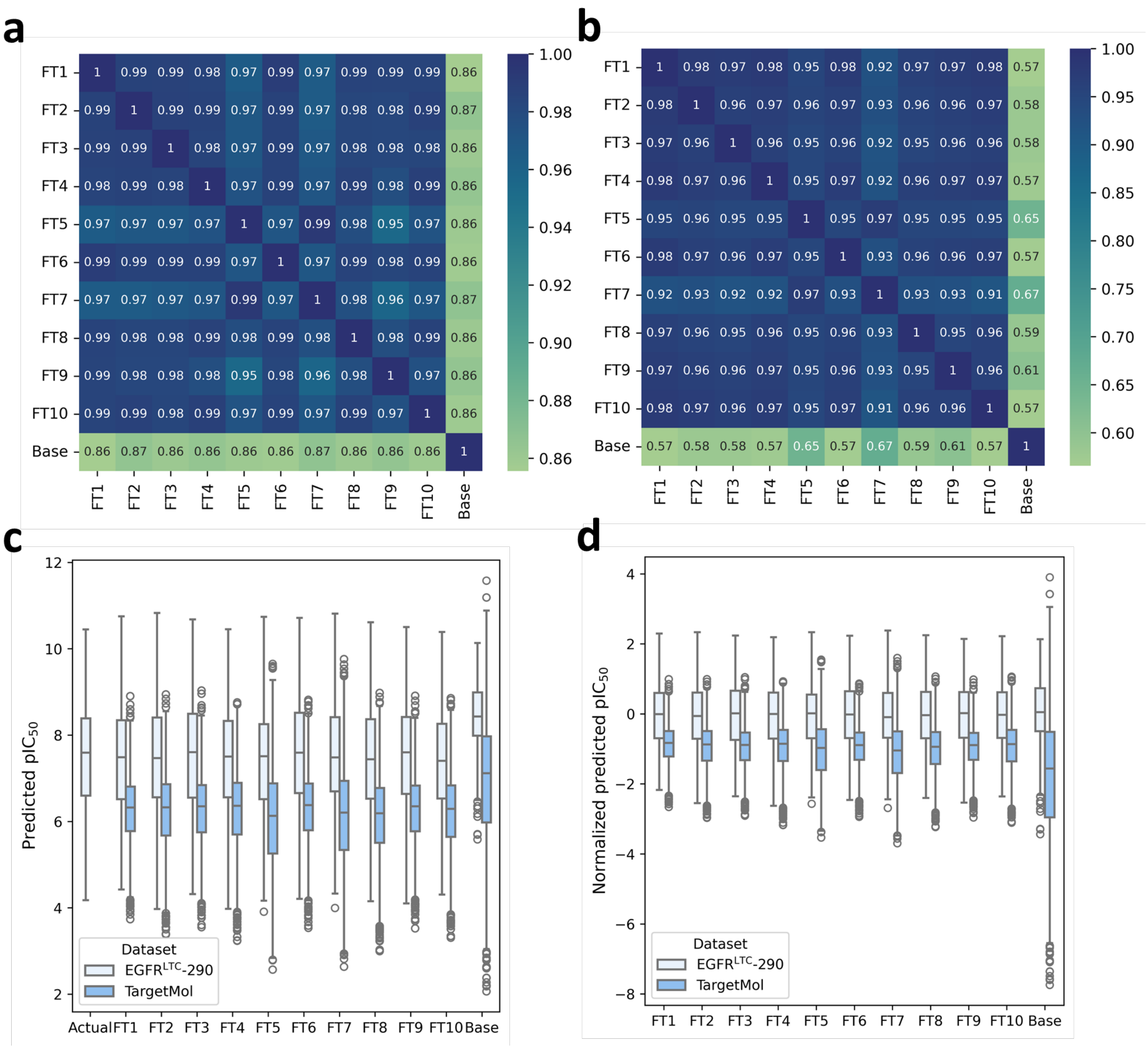


**Figure S9. Comparison of predicted affinities from the base model (Model B) and fine-tuned models (FT1 to FT10) on EGFR$^{LTC}$-290 dataset and TargetMol library. (a)** Correlation matrix of predicted affinities between different models on the EGFR$^{LTC}$-290 dataset. **(b)** Correlation matrix of predicted affinities between different models on the TargetMol library. **(c)** Distribution of predicted affinities from the base model and fine-tuned models on the $EGFR^{LTC}$-290 dataset and TargetMol library. The first column is the distribution of measured $pIC_{50}$ in $EGFR^{LTC}$-290. **(d)** Normalized distribution of predicted affinities on the $EGFR^{LTC}$-290 dataset. This normalization helps to make the predictions from different models more comparable.

**Figure S9a** displays the correlations between the predicted affinities of the 11 models employed in the screening on the $EGFR^{LTC}$-290 dataset, which includes the base model (Model B) and the ten fine-tuned models (FT1 to FT10). The results suggest a high consistency between the predictions made by the ten fine-tuned models, with an average correlation coefficient of 0.98. However, predictions between the base model and the fine-tuned models sometimes vary, with an average correlation coefficient of 0.86. Similar phenomena were also observed with the TargetMol library (**Figure S9b**).

We observed that the distribution of predicted affinity of the base model differs substantially from the distribution of predictions from the fine-tuned models (**Figure S9c**). The base model has a higher average but a smaller deviation on the $EGFR^{LTC}$-290 dataset, while it has a higher average but larger deviation on the TargetMol library. This discrepancy might be due to the fact that the base model was trained to predict $pK_d$, whereas the measure of affinity used in this study is represented by $pIC_{50}$, which is typically lower than $pK_d$.

To make the predictions more comparable, we normalized them to the uniform distribution of predictions on the $EGFR^{LTC}$-290 dataset (**Figure S9d**). In addition to considering affinity, we also took into account other factors when making our final selection (details are in Methods).

In **Figure S10**, we present visualizations of the chemical space comparing the TargetMol library with the $EGFR^{LTC}$-290 dataset. These visualizations of PCA and t-SNE plots reveal that compounds in the TargetMol library possess compounds significantly different from those in $EGFR^{LTC}$-290, particularly noticeable at the bottom of these plots. Despite these chemical differences, these compounds were not predicted to be highly effective inhibitors (**Figure S10d**). Further analysis shows the distinct chemical spaces occupied by compounds C1, C5, C7, and C11 as compared to the existing $EGFR^{LTC}$-290 inhibitors (**Figure S10a**). In contrast, C4 and C10 are observed to be closer to other $EGFR^{LTC}$ inhibitors in these 2D plots. This proximity is attributed to the shared aniline-pyrimidine scaffold common to C4, C10, and many existing $EGFR^{LTC}$ inhibitors. Additionally, **Figure S11** highlights the precision of the fine-tuned model in predicting the IC50 values for compounds C1, C10, and C11. Notably, this precise prediction is achieved even though C1 and C11 differ significantly from the compounds in the $EGFR^{LTC}$-290.

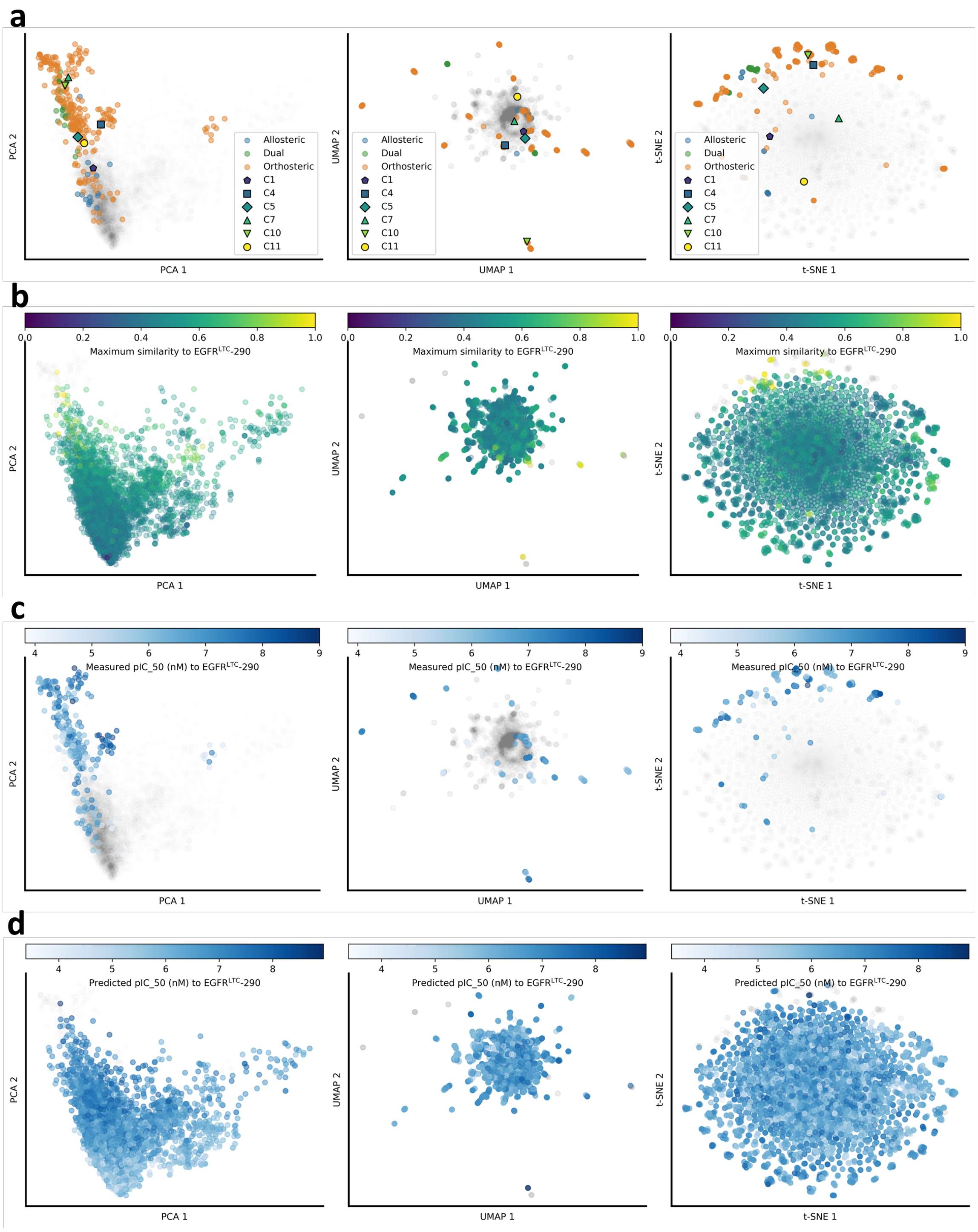


**Figure S10. Visualization of chemical space for EGFR$^{LTC}$-290, TargetMol library, and selected inhibitors.** Displayed are PCA, UMAP, and t-SNE projections, from left to right, based on Morgan fingerprints[8]. **(a)** EGFR$^{LTC}$-290 inhibitors are coloured in blue, green, and orange to represent allosteric, dual, and orthosteric types, respectively. TargetMol library compounds are shown in grey, with newly identified inhibitors in distinct colors. **(b)** EGFR$^{LTC}$-290 inhibitors are shown in grey, while TargetMol library compounds are shaded according to their highest Tanimoto similarity to EGFR$^{LTC}$-290 compounds, transitioning from purple (low similarity) to yellow (high similarity), based on topological fingerprints in RDKit[11]. **(c)** TargetMol library compounds are shown in grey, with EGFR$^{LTC}$-290 compounds coloured by their measured $pIC_{50}$ values, ranging from white (low) to blue (high). (d) EGFR$^{LTC}$-290 compounds are shown in grey, with TargetMol library compounds shaded according to their predicted $pIC_{50}$ values (average predictions from models FT1 to FT10), ranging from white (low) to blue (high).

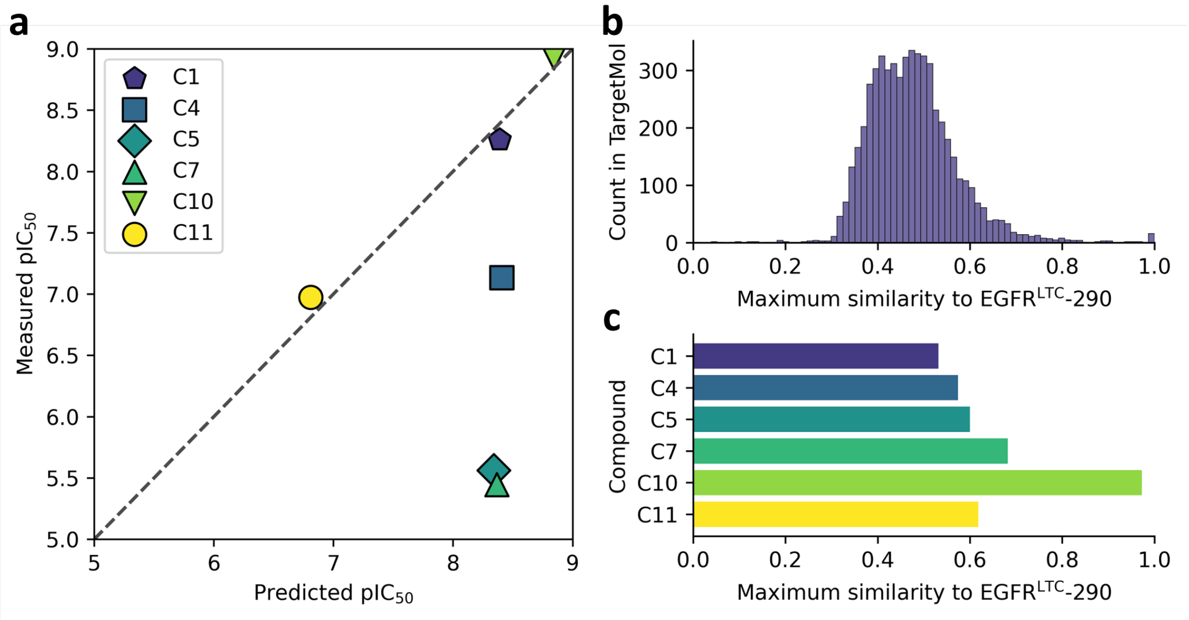


**Figure S11. Analysis of identified EGFR$^{LTC}$ inhibitors. (a)** Scatter plot comparing the predicted versus measured $pIC_{50}$ values for identified inhibitors. Each point is coloured to correspond to a specific inhibitor, plotted against the dashed line representing perfect prediction accuracy. **(b)** Histogram showing the distribution of maximum Tanimoto similarities for compounds in the TargetMol library to the EGFR$^{LTC}$-290 set **(c)** Bar graph illustrating the maximum Tanimoto similarity of each inhibitor to the EGFR$^{LTC}$-290 set. Similarity calculations are based on topological fingerprints in RDKit[11].

### Supplementary References


1 Jumper, J. *et al.* Highly accurate protein structure prediction with AlphaFold. *Nature* **596**, 583-589 (2021).

2 Liu, S. *et al.* Pre-training molecular graph representation with 3D geometry. *arXiv* 2110.07728 (2021).

3 Liu, Z. *et al.* PDB-wide collection of binding data: Current status of the PDBbind database. *Bioinformatics* **31**, 405-412 (2015).

4 Lu, W. *et al.* Tankbind: Trigonometry-aware neural networks for drug-protein binding structure prediction. *Adv. Neural Inf. Process. Syst.* **35**, 7236-7249 (2022).

5 Li, S. *et al.* Monn: A multi-objective neural network for predicting compound-protein interactions and affinities. *Cell Syst.* **10**, 308-322. e311 (2020).

6 Kyro, G. W., Brent, R. I. & Batista, V. S. Hac-Net: A hybrid attention-based convolutional neural network for highly accurate protein–ligand binding affinity prediction. *J. Chem. Inf. Model.* **63**, 1947-1960 (2023).

7 Van Der Maaten, L. Accelerating t-SNE using tree-based algorithms. *J. Mach. Learn. Res.* **15**, 3221-3245 (2014).

8 Rogers, D. & Hahn, M. Extended-connectivity fingerprints. *J. Chem. Inf. Model.* **50**, 742-754 (2010).

9 Kyte, J. & Doolittle, R. F. A simple method for displaying the hydropathic character of a protein. *J. Mol. Biol.* **157**, 105-132 (1982).

10 Zamyatnin, A. Protein volume in solution. *Prog. Biophys. Mol. Biol.* **24**, 107-123 (1972).

11 Landrum, G. RDKit: Open-source cheminformatics. (2006).

12 Cock, P. J. *et al.* Biopython: Freely available python tools for computational molecular biology and bioinformatics. *Bioinformatics* **25**, 1422 (2009).

13 Ba, J. L., Kiros, J. R. & Hinton, G. E. Layer normalization. *arXiv* 1607.06450 (2016).

14 Srivastava, N., Hinton, G., Krizhevsky, A., Sutskever, I. & Salakhutdinov, R. Dropout: A simple way to prevent neural networks from overfitting. *J. Mach. Learn. Res.* **15**, 1929-1958 (2014).

15 Vaswani, A. *et al.* Attention is all you need. *Adv. Neural Inf. Process. Syst.* **30** (2017).

16 He, K., Zhang, X., Ren, S. & Sun, J. Deep residual learning for image recognition. *Proc. IEEE Comput. Soc. Conf. Comput. Vis. Pattern Recognit.*, 770-778 (2016).

17 Kingma, D. P. & Ba, J. Adam: A method for stochastic optimization. *arXiv* 1412.6980 (2014).

18 Mistry, J. *et al.* Pfam: The protein families database in 2021. *Nucleic Acids Res.* **49**, D412-D419 (2021).
